# *Salmonella* Typhimurium uses anaerobic respiration to overcome propionate-mediated colonization resistance

**DOI:** 10.1101/2021.05.25.445690

**Authors:** Catherine D. Shelton, Woongjae Yoo, Nicolas G. Shealy, Teresa P. Torres, Jacob K. Zieba, M. Wade Calcutt, Nora J. Foegeding, Dajeong Kim, Jinshil Kim, Sangryeol Ryu, Mariana X. Byndloss

## Abstract

The gut microbiota benefits the host by limiting enteric pathogen expansion (colonization resistance) partially via the production of inhibitory metabolites. Propionate, a short-chain fatty acid produced by microbiota members, is proposed to mediate colonization resistance against *Salmonella enterica* serovar Typhimurium (*S.* Tm). Here, we show that *S.* Tm overcomes the inhibitory effects of propionate by using it as a carbon source for anaerobic respiration. We determined that propionate metabolism provides an inflammation-dependent colonization advantage to *S.* Tm during infection. Such benefit was abolished in the intestinal lumen of *Salmonella*-infected germ-free mice. Interestingly, S. Tm propionate-mediated intestinal expansion was restored when germ-free mice were monocolonized with *Bacteroides thetaiotaomicron (B. theta),* a prominent propionate producer in the gut, but not when mice were monocolonized with a propionate production-deficient *B. theta* strain. Taken together, our results reveal a novel strategy used by *S.* Tm to mitigate colonization resistance by metabolizing microbiota-derived propionate.

## Introduction

The intestines are occupied by a complex microbial community, the gut microbiota, mainly composed of obligate anaerobic bacteria. By residing in the gut, the microbiota contributes to host health through nutrient production (1, 2), immune education (3, Reviewed in 4), and protection against enteric pathogens (colonization resistance) (5, 6). Colonization resistance is accomplished through diverse mechanisms (7). For instance, the gut microbiota can indirectly inhibit pathogen expansion by activating the host’s immune response or by enhancing the intestinal mucosal barrier (8, 9, 10). On the other hand, direct antagonism of enteric pathogens by the gut microbiota is achieved through niche competition or the production of inhibitory molecules (11, 12).

A proposed microbiota-derived metabolite that mediates colonization resistance is propionate (13, 14). Propionate, an abundant short-chain fatty acid (15), is generated by the fermentation of sugars by anaerobic bacteria, specifically members of the *Bacteroides* genus (16). As a predicted component of colonization resistance, propionate has previously been studied for its specific role in inhibiting *Salmonella enterica* serovar Typhimurium (*S.* Tm). The complete mechanism by which propionate exerts its toxic effect on *S.* Tm remains unknown. Initially, propionate was shown to inhibit *S.* Tm by generating toxic by-products produced during propionate catabolism (14). Propionate was recently shown to acidify the intracellular space of *S.* Tm and significantly reduce the *S*. Tm’s growth rate (13). However, the mechanisms employed by enteric pathogens to overpower propionate-mediated colonization resistance, a key step for successful gut colonization, remain largely unknown.

*S.* Tm has evolved several mechanisms to overcome colonization resistance. Upon infection, *S.* Tm invades the intestinal epithelium and activates the host’s innate immune system and inflammatory response (17). As a result, the host produces reactive nitrogen species (RNS), specifically nitric oxide, to inhibit the growth of the pathogen (18). Host-generated nitric oxide can react with other compounds in the intestinal lumen to generate nitrate (19, 20). Interestingly, *S*. Tm can take advantage of the nitrate generated by the host immune response by using it as an alternative electron acceptor to fuel anaerobic respiration (21). By performing anaerobic respiration, *S.* Tm can outgrow the resident microbiota whose metabolism relies on fermentation (22, 23). In addition to the energetic benefits of anaerobic respiration compared to fermentation, *S.* Tm’s metabolic adaption in the inflamed gut enables the pathogen to access new nutrient niches and metabolize novel carbon sources (24, 25).

The carbon sources utilized by *S.* Tm during anaerobic respiration remain largely uncharacterized. Interestingly, is it possible that *S*. Tm may use propionate as a carbon source during infection, as this pathogen possesses the machinery necessary for propionate catabolism (26, 27). Specifically, the *prpBCDE* operon encodes the enzymes required to convert propionate into pyruvate through the 2-methylcitrate cycle (27). Furthermore, genes in the *prp* operon are nonfunctional in extraintestinal serovars of *S.* Tm, raising the possibility that propionate metabolism is required for successful *S.* Tm colonization in the inflamed intestinal lumen (28). In this study, we show that the ability of *S*. Tm to metabolize propionate relies on nitrate-dependent anaerobic respiration. We then use conventional and germ-free mouse models of *S*. Tm gastroenteritis to demonstrate that *S*. Tm uses inflammation-dependent anaerobic respiration to overcome propionate-mediated colonization resistance.

## Results

### Propionate supports *S.* Tm growth during anaerobic respiration *in vitro*

Previous investigations into the function of the *prp* operon have focused on the use of propionate as a carbon source under aerobic conditions (14, 29) (**Figure 1A and S1A**). However, as an enteric pathogen, *S*. Tm encounters propionate in the intestinal lumen, which is mainly anaerobic (30). Therefore, we sought to investigate the ability of *S*. Tm to metabolize propionate under conditions relevant to intestinal disease. We first determined if propionate could support *S.* Tm growth *in vitro* if alternative electron acceptors (i.e., DMSO, TMAO, fumarate, tetrathionate, nitrate) generated during *S.* Tm-induced inflammation were available (20, 28, 31) (**Figure 1B**). Interestingly, we observed that propionate significantly increased *S*. Tm growth only when nitrate was added to the media (**Figure 1B**). Propionate did not increase *S.* Tm growth when the other alternative electron acceptors were added (**Figure 1B**). We next tested if a range of propionate concentrations (5 – 50 mM), physiologically relevant for mice and humans (15, 16), supported *S.* Tm growth in the presence or absence of nitrate. *S.* Tm was able to grow in increasing concentrations of propionate when nitrate was present in the media (**Figure 1C**), suggesting that *S*. Tm may be able to metabolize high levels of propionate during infection. Notably, *S.* Tm could not ferment propionate as no growth is observed in the absence of nitrate (**Figure 1C**).

**Figure 1.**
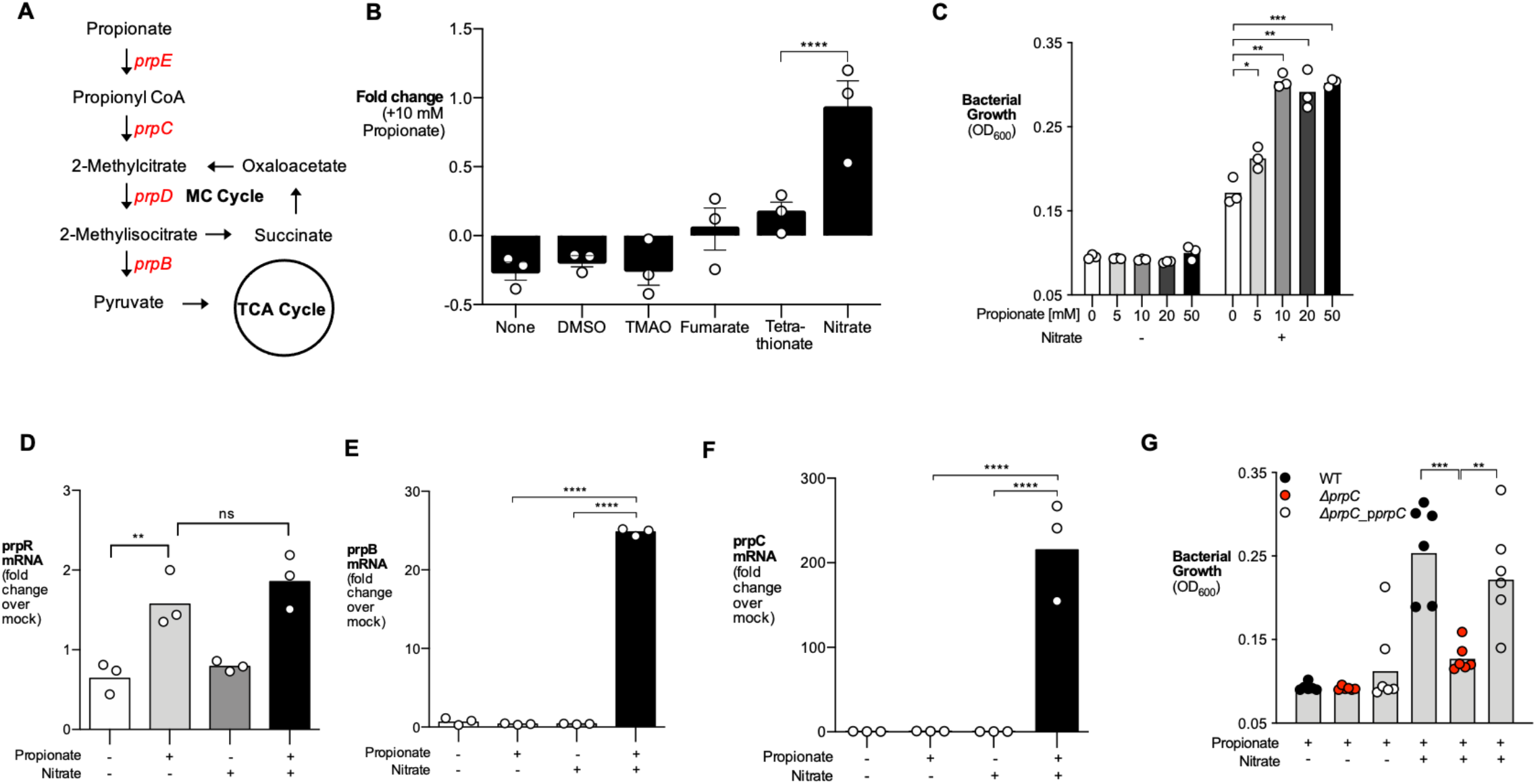
Propionate fuels *S.* Tm growth in the presence of nitrate *in vitro.* (A) Simplified model of propionate catabolism in *S.* Tm. Genes in the *prpBCDE* operon (red) metabolize propionate into pyruvate. *prpE,* propionyl-CoA synthase; *prpC*, methylcitrate synthase; *prpD*, methylcitrate dehydratase; *prpB*, 2-methylisocitrate lyase. (B) NCE minimal media containing 40 mM of an alternative electron acceptor alone or 40 mM alternative electron acceptor + 10 mM propionate was inoculated with *S.* Tm and grown anaerobically for 24 hours. Fold change calculated by comparing growth of alternative electron acceptor + 10 mM propionate to growth with alternative electron acceptor alone. (C) NCE minimal media containing increasing propionate concentration with or without 40 mM nitrate was inoculated with *S.* Tm. OD_600_ of *S.* Tm was measured after 24 hours of anaerobic growth. (D – F) Relative transcription of *prpR* (D), *prpB* (E), *prpC* (F) in NCE minimal media supplemented with propionate, nitrate, or both propionate and nitrate was determined by qRT-PCR. Transcription of target genes was normalized to *gyrB* rRNA. (G) NCE minimal media containing propionate or propionate and nitrate was inoculated with wildtype *S.* Tm, Δ*prpC,* or a complemented strain of Δ*prpC* (Δ*prpC_*p*prpC).* Media was supplemented with 200 μM Isopropyl β-D-1-thiogalactopyranoside (IPTG) to induce expression of *prpC* in the complemented mutant strain. OD_600_ of each strain was measured after 24 hours of anaerobic growth. Each dot represents one biological replicate. Bars represent the geometric mean. *, p < 0.05; **, p < 0.01; ***, p < 0.001; ****, p < 0.0001; ns, not statistically significant. See also Figure S1.

To determine whether anaerobic respiration affects expression of the *prpBCDE* operon, we measured changes in *S.* Tm gene transcription when propionate, nitrate, or propionate and nitrate were available. Propionate alone induced expression of *prpR,* the transcriptional activator of the *prpBCDE* operon (**Figure 1D**). However, addition of nitrate (nitrate + propionate media) was necessary to increase expression of the propionate utilization genes *prpB* and *prpC* (**Figure 1E and 1F**), supporting the specific role of nitrate in propionate catabolism. We next confirmed that the *prpBCDE* operon was necessary for *S.* Tm growth in the presence of propionate and nitrate. Deletion of the *prpBCDE* operon or *prpC* in *S.* Tm blunted the pathogen’s growth on propionate under anaerobic respiration conditions (**Figure 1G, S1B, S1C**). However, no defects in growth were observed when mutants were given glucose or glycerol as a carbon source (**Figure S1D and S1E**). Growth on propionate could be restored in Δ*prpC* by reintroducing the *prpC* gene by plasmid complementation (**Figure 1G**). These experiments reveal that nitrate respiration supports propionate catabolism *in vitro* through the *prpBCDE* operon.

### Nitrate respiration allows *S*. Tm to overcome the inhibitory effects of propionate under low pH

The ability of *S*. Tm to use propionate as a carbon source under anaerobic respiration conditions may be confounded by the acidifying effects of this short-chain fatty acid (13, 32), particularly in the low pH of the intestines (ranging from pH 7.4 to 5.7) (33). Thus, we determined how changes in pH impact the growth of *S.* Tm with propionate and nitrate *in vitro*. Notably, *S.* Tm grew significantly more at pH 7.0, pH 6.5, and pH 6.0 when propionate and nitrate were present than with propionate alone (**Figure 2A-C**). Wildtype *S.* Tm also grew significantly better than Δ*prpC*, which failed to grow when given propionate and nitrate at all pH values tested (**Figure 2A-C**). The growth of wildtype *S.* Tm at pH 6.0 was reduced compared to growth at pH 6.5 and pH 7.0 (**Figure 2C**). However, decreased growth is observed when *S.* Tm is grown with glycerol and nitrate at pH 6.0, suggesting that reduced growth at pH 6.0 is not specific to propionate metabolism (**Figure S2A and S2B**).

**Figure 2.**
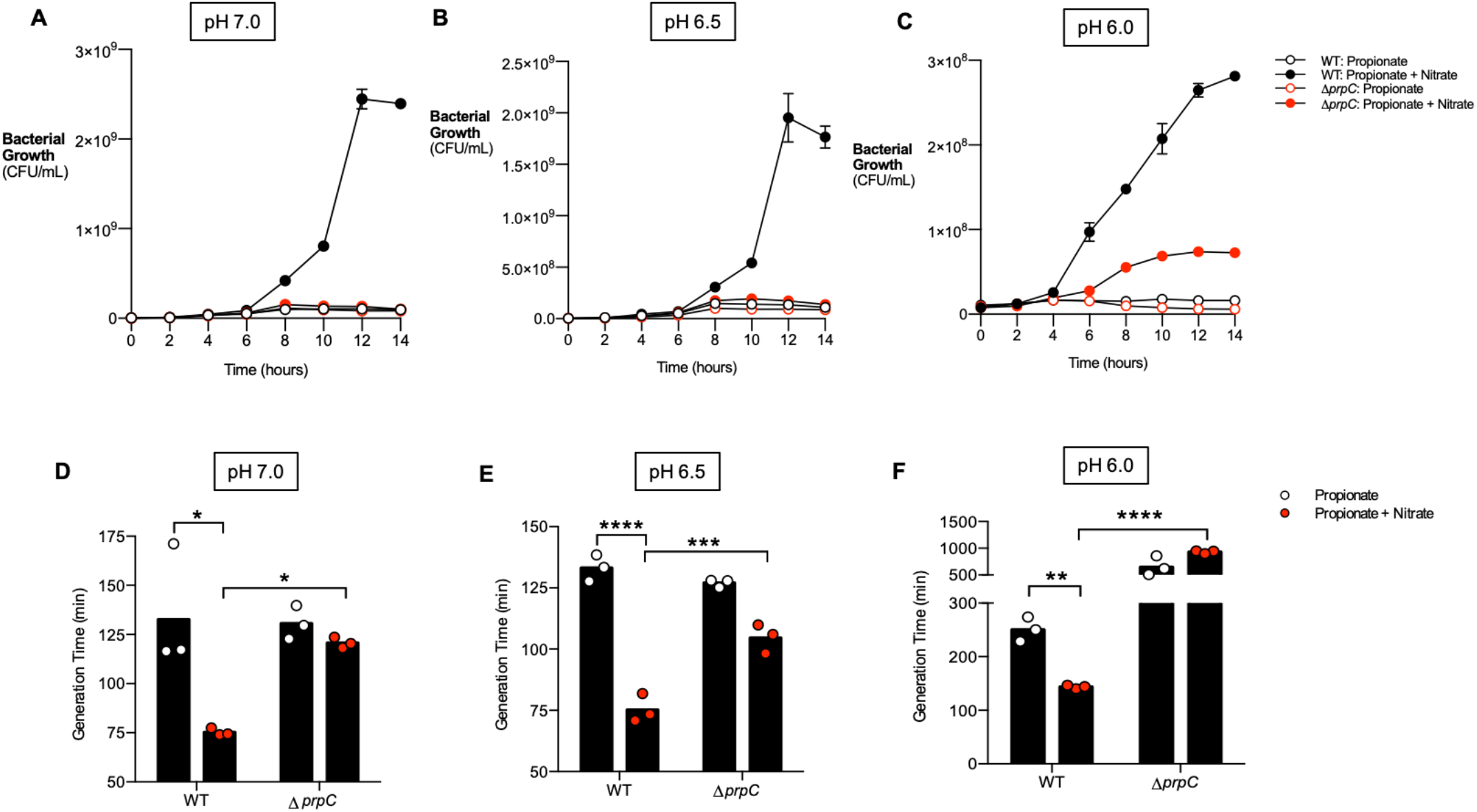
Nitrate prevents inhibitory effects of propionate on *S.* Tm *in vitro*. (A-C) NCE minimal media was adjusted to pH 7.0 (A), pH 6.5 (B), and pH 6.0 (C). Media was supplemented with 10 mM propionate or 10 mM propionate + 40 mM nitrate and inoculated with wildtype *S.* Tm or Δ*prpC.* The culture was grown anaerobically for 14 hours, and samples were taken every two hours to plate for CFUs. N=4. (D-F) Cultures grown in A-C were used to calculate the generation time of wildtype *S.* Tm or Δ*prpC* at pH 7.0 (D), pH 6.5 (E), pH 6.0 (F). Each dot represents one biological replicate. Bars represent the geometric mean. *, p < 0.05; **, p < 0.01; ****, p < 0.0001. See also Figure S2.

Previous research revealed that propionate mediated colonization resistance by decreasing the growth rate of *S.* Tm (13). To address if nitrate prevented this effect of propionate, the generation time of *S.* Tm at pH 7.0, 6.5, and 6.0 was calculated when *S*. Tm was grown in the presence of propionate and nitrate or propionate alone. The addition of nitrate significantly decreased the generation time of wildtype *S.* Tm at all pH values tested (**Figure 2D-F**). Furthermore, the generation time of wildtype *S.* Tm was markedly shorter than Δ*prpC* at each pH tested when propionate and nitrate were present (**Figure 2D-F**). Although nitrate led to a significant reduction in generation time for Δ*prpC* at pH 6.5, no difference in generation time was observed at pH 7.0 and pH 6.0 when both nitrate and propionate were available (**Figure 2D-F**). These data show that nitrate can mitigate the inhibitory effects of propionate on *S.* Tm growth by promoting propionate metabolism through the *prpBCDE* operon.

### Propionate metabolism provides a growth advantage to *S.* Tm in the inflamed gut

After discovering that *S.* Tm can metabolize propionate under anaerobic respiration conditions *in vitro*, we next investigated whether propionate catabolism provided *S*. Tm with a colonization advantage *in vivo*. We infected C57BL/6 mice, pretreated with streptomycin, with an equal mixture of wildtype *S.* Tm and Δ*prpC*. Four days after infection, the bacterial load of each strain was determined by plating the colon contents on selective media, and the ratio of *S*. Tm wildtype and Δ*prpC* (competitive index) was calculated. To determine the role of propionate catabolism in *S*. Tm colonization of systemic sites, we also assessed *S*. Tm wildtype vs. *ΔprpC* mutant competitive index in spleen and liver sampled from infected mice. Interestingly, we observed a significant competitive advantage for WT *S.* Tm over Δ*prpC* in the colon contents, but not in the liver or spleen (**Figure 3A).** These experiments reveal that propionate metabolism benefits *S.* Tm during infection specifically in the gastrointestinal tract.

**Figure 3.**
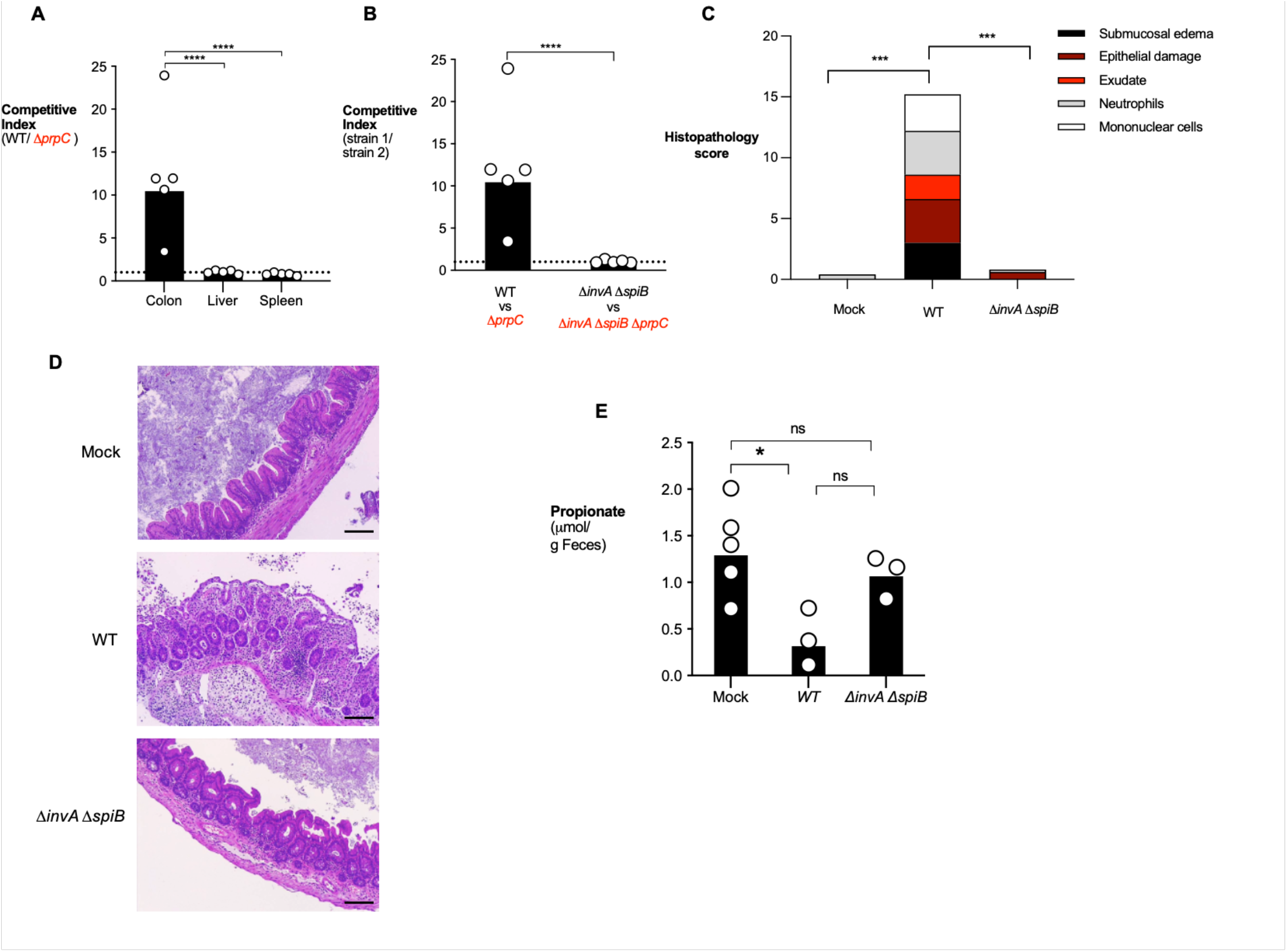
Propionate utilization confers an advantage to *S.* Tm in an inflammation-dependent mechanism. (A) Streptomycin-pretreated C57BL/6 mice were inoculated with an equal mixture of wildtype *S.* Tm and Δ*prpC.* The competitive index in the colon content and homogenized samples from the liver or spleen was determined four days after infection. (B) Streptomycin-pretreated C57BL/6 mice were inoculated with an equal mixture of the indicated S. Tm strains. The competitive index in the colonic content was determined four days after infection. (C) Combined histopathology score of pathological lesions in the cecum of mice from (B). (D) Representative images of Hematoxylin and Eosin-stained cecal tissue of mice from (C). Scale bar equals 200 μm. (E) Propionate concentration in the cecal content was determined by liquid chromatography/ mass spectrometry (LC/MS) two days after infection. Each dot represents one animal. Bars represent the geometric mean. *, p < 0.05; **, p < 0.01; ***, p < 0.001; ****, p < 0.0001.

Next, we investigated whether inflammation was required for propionate metabolism to confer an advantage to *S.* Tm. C57BL/6 mice, pretreated with streptomycin, developed significant intestinal inflammation characterized by edema, epithelial damage, infiltration of inflammatory cells in the submucosa and exudate in the intestinal lumen four days after *S*. Tm infection (**Fig. 3C, D**). *S.* Tm uses two type III secretion systems (T3SS) to invade the intestinal epithelium and perform intracellular replication (17). A mutant strain (Δ*invA* Δ*spiB*) of *S.* Tm is defective in both T3SS and does not cause inflammation in a mouse model (34, 35). Thus, we constructed an Δ*prpC* mutant in the *S.* Tm inflammation-deficient background (Δ*invA* Δ*spiB* Δ*prpC*) and then infected streptomycin pretreated C57BL/6 mice with an equal mixture of Δ*invA* Δ*spiB* and Δ*invA* Δ*spiB* Δ*prpC*. The competitive index was determined four days after infection. In contrast to the competitive advantage observed for wildtype over Δ*prpC,* no competitive advantage was observed for Δ*invA* Δ*spiB* over Δ*invA* Δ*spiB* Δ*prpC,* revealing that inflammation is required for propionate metabolism to be advantageous to *S.* Tm (**Figure 3B**). Histopathology analysis confirmed that infection with *S*. Tm ΔinvA ΔspiB did not induce intestinal inflammation (**Fig. 3C, D**). The concentration of propionate was measured in the feces of mice infected with wildtype or Δ*invA* Δ*spiB* and no significant differences were observed (**Figure 3E**), indicating that the lack of an advantage of Δ*invA* Δ*spiB* over Δ*invA* Δ*spiB* Δ*prpC* was not due to decreased levels of propionate in Δ*invA* Δ*spiB*-infected mice.

### Nitrate respiration is required for *S.* Tm to benefit from propionate metabolism *in vivo*

We observed *in vitro* that the only alternative electron acceptor that can support *S.* Tm growth on propionate is nitrate (**Fig. 1**). To perform anaerobic respiration using nitrate as an alternative electron acceptor, *S.* Tm relies on three nitrate reductases, *narGHI, narZYV, napABC* (36). Indeed, a Δ*napA* Δ*narZ* Δ*narG* mutant strain of *S.* Tm is unable to perform nitrate-dependent anaerobic respiration (22, 36). To investigate if nitrate respiration is required for propionate metabolism to be advantageous *in vivo*, we infected streptomycin pretreated C57BL/6 mice with an equal mixture of *S*. Tm Δ*napA* Δ*narZ* Δ*narG* and *S*. Tm Δ*napA* Δ*narZ* Δ*narG* Δ*prpC* and measured the competitive index four days after infection. In contrast to the competitive advantage observed for wildtype over Δ*prpC,* no advantage was observed for Δ*napA* Δ*narZ* Δ*narG* over Δ*napA* Δ*narZ* Δ*narG* Δ*prpC* (**Figure 4A**), suggesting that nitrate respiration is required for propionate metabolism to benefit *S.* Tm *in vivo*.

**Figure 4.**
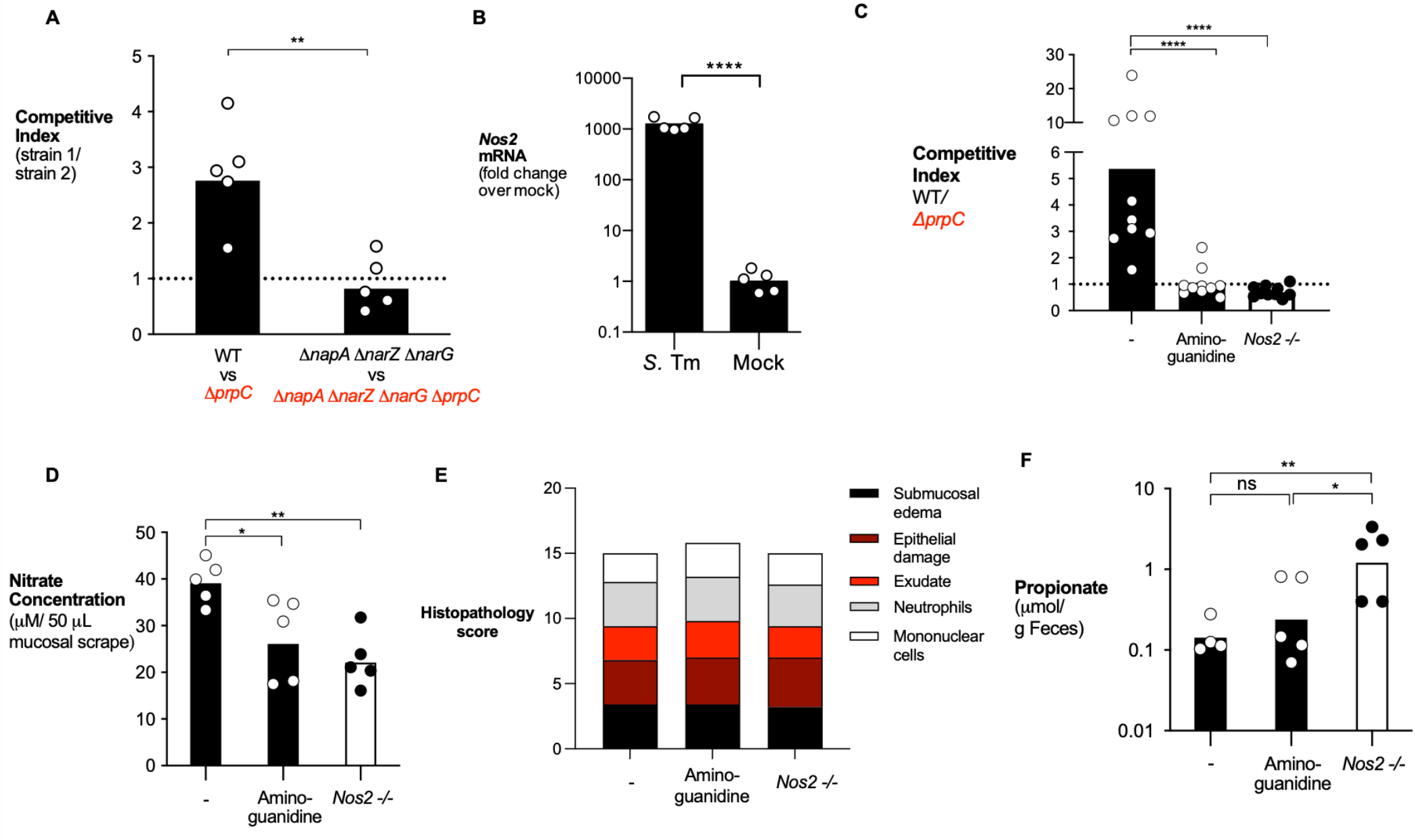
Inflammation generated nitrate required for propionate catabolism by *S.* Tm. (A) Streptomycin-pretreated C57BL/6 mice were inoculated with an equal mixture of the indicated S. Tm strains. The competitive index in the colon content was determined four days after infection. (B) Streptomycin-pretreated C57BL/6 mice were inoculated with *S.* Tm or mock-treated. mRNA levels of *Nos2* were measured in the cecal mucosa 4 days post-infection and normalized to β-actin mRNA levels. (C) Streptomycin-pretreated C57BL/6 wildtype mice and Nos2-deficient mice were inoculated with an equal mixture of the wildtype *S*. Tm and Δ*prpC* mutant. One group was treated with aminoguanidine as indicated. The competitive index in the cecal content was determined four days post-infection. (D) Nitrate concentration in the colonic mucus layer was determined in wildtype mice, wildtype mice treated with aminoguanidine, and Nos2-deficient mice by a modified Griess assay. (E) Combined histopathology score of pathological lesions in the cecum of mice from (C). (F) Propionate concentration in the cecal content was determined by liquid chromatography/ mass spectrometry (LC/MS) two days after infection. Each dot represents one animal. Bars represent the geometric mean. *, p < 0.05; **, p < 0.01; ***, p < 0.001; ****, p < 0.0001.

As an alternative approach, we investigated if propionate metabolism was advantageous to *S.* Tm if the availability of inflammation-derived nitrate was decreased. During *Salmonella* infection, inflammatory monocytes and intestinal epithelial cells upregulate inducible nitric oxide synthase (iNOS) encoded by *Nos2*, leading to the production of nitric oxide (22, 23) (**Figure 4B**). Nitric oxide reacts with superoxide radicals to form peroxynitrate, which can then decompose into nitrate (31). Therefore, we repeated the competitive infection assays in mice treated with the iNOS inhibitor aminoguanidine (31), and in iNOS (*Nos2*)-deficient mice. In mock-treated wildtype mice, we observed a competitive advantage for wildtype *S.* Tm over Δ*prpC* (**Figure 4C**). However, this advantage was abrogated both in mice treated with aminoguanidine and in *Nos2*-deficient mice (**Figure 4C**). Nitrate measurements from colonic and cecal mucosa revealed a significant decrease in nitrate levels in mice treated with aminoguanidine and in *Nos2-*deficient mice (**Figure 4D**). Despite differences in nitrate levels, inflammation was similar between the three treatment groups (**Figure 4E**). Propionate levels were similar in the feces of aminoguanidine-treated mice and were elevated in the feces of Nos2 −/− mice compared to mock-treated wildtype mice (**Figure 4F**), confirming that lack of a competitive advantage of wildtype *S.* Tm over *ΔprpC* in aminoguanidine-treated or in Nos2 −/− mice was not due to decreased propionate or differences in inflammation between experimental groups. Together, these data reveal that the host-derived nitrate supports propionate utilization by *S.* Tm during pathogen-induced gastroenteritis.

### Microbiota-derived propionate provides a growth advantage to *S.* Tm in the presence of nitrate

While we predict that the advantage of wildtype *S.* Tm over Δ*prpC* (**Figure 3**) is due to the ability of wildtype *S.* Tm to overcome propionate-mediated colonization resistance, it is possible that other microbiota-derived metabolites may contribute to this phenotype. Therefore, we sought to examine the advantage of wildtype versus Δ*prpC* using an *in vitro* approach in which propionate levels can be controlled. As a source of propionate, we used *Bacteroides thetaiotaomicron,* a representative of the *Bacteroides* genus and a predominant propionate producer in the gut (37). As a negative control, we also cultured *B. theta* BT1686-89, an isogenic propionate production-deficient *B. theta* mutant strain (38). *B. theta* BT1686-89 was grown anaerobically in mucin-broth for four days to adjust for a growth defect, while wildtype *B. theta* was grown for two days (**Figure S3A and S3B**). Growth of wildtype *B. theta* led to an accumulation of propionate in the media while *B. theta* BT1686-89 supernatant contained significantly less propionate (**Figure 5A**). Then, to determine whether microbiota-derived propionate provides a growth advantage to *S.* Tm, we cultured wildtype *S.* Tm and *S.* Tm Δ*prpC* in supernatant from wildtype *B. theta* or *B. theta* BT1686-89, and *B. theta* supernatants were either left untreated or supplemented with nitrate (**Figure 5B-C**). In the absence of nitrate, no differences in growth were observed between wildtype *S*. Tm or Δ*prpC* cultured in supernatant from either wildtype *B. theta* or *B. theta* BT1686-89 (**Figure 5A**). However, addition of nitrate enabled wildtype *S.* Tm to grow to significantly higher levels than Δ*prpC* in supernatant from wildtype *B. theta*, but not in supernatant from *B. theta* BT1686-89 (**Figure 5B-C**). Restoring propionate levels in supernatant of *B. theta* BT1686-89 (**Figure 5A**) rescued wildtype *S.* Tm’s ability to grow significantly more than Δ*prpC* in the presence of nitrate (**Figure 5B-C**). These results indicate that the growth advantage of wildtype *S.* Tm over Δ*prpC* is specific to propionate and not due to other *B. theta*-derived metabolites. Moreover, these results demonstrate that microbe-derived propionate can fuel *S.* Tm growth in the presence of nitrate in a *prpBCDE-*dependent manner.

**Figure 5.**
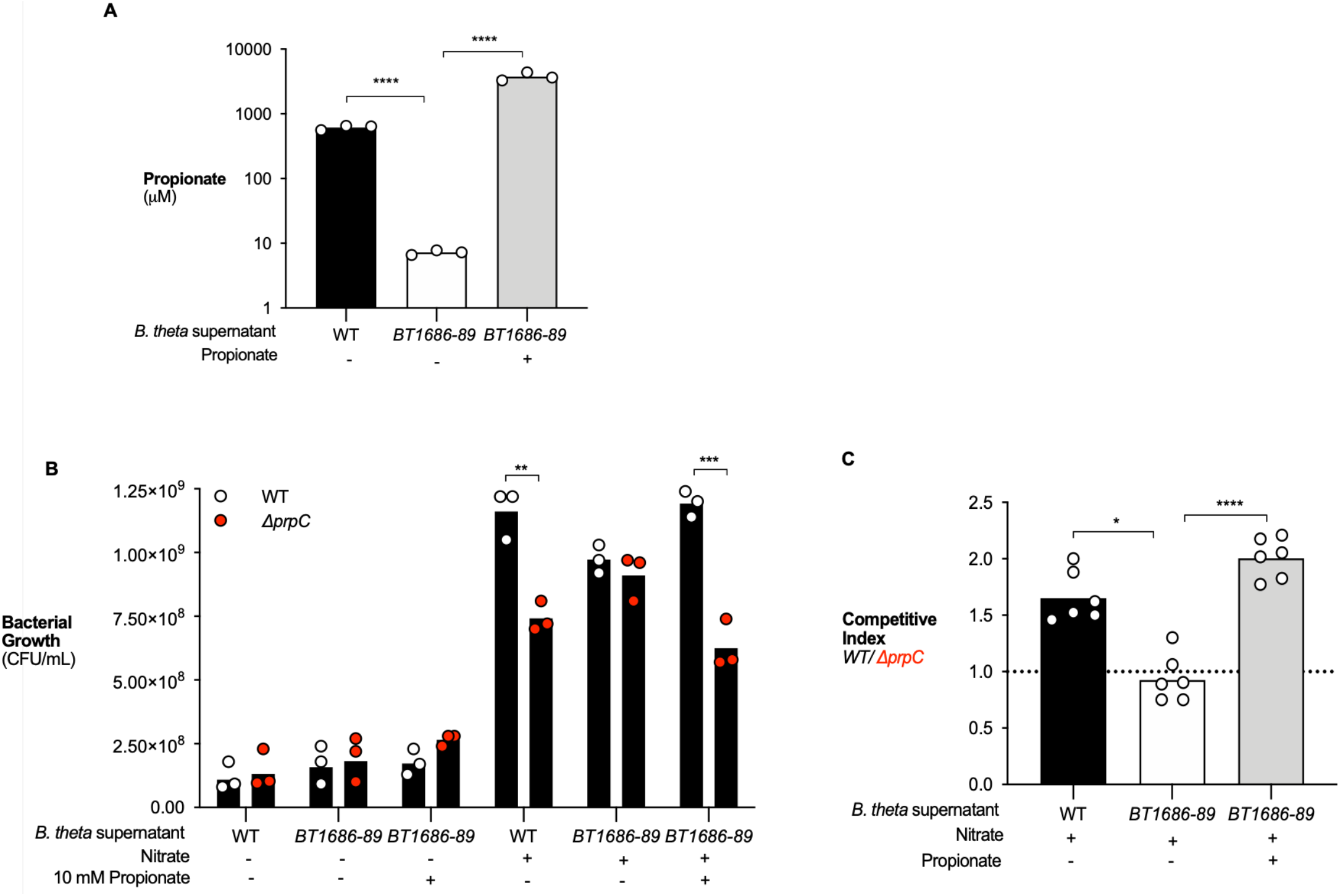
*Bacteroides* produced propionate is a carbon source for *S.* Tm if nitrate is present. (A-C) Mucin broth was inoculated with wildtype *B. thetaiotaomicron (WT*) *or B. thetaiotaomicron* BT1686-89 (BT1686-89*). B. theta* BT1686-89 was cultured anaerobically for 4 days and wildtype *B. theta* was cultured for 2 days. (A) Propionate concentration in the digested mucin broth from WT or BT1686-89 *B. theta* culture (supplemented or not with propionate) was determined by liquid chromatography/ mass spectrometry (LC/MS). (B) Filter-sterilized wildtype *B. theta* or *B. theta* BT1686-89*-digested* mucin broth was inoculated with an equal mixture of wildtype *S.* Tm and Δ*prpC*. 40 mM nitrate and 10 mM propionate were added where indicated. Cultures were plated after 16 hours of anaerobic growth. (C) Competitive index was calculated from CFU counts in (A). Nitrate and propionate were added as indicated. Each dot represents one biological replicate. Bars represent the geometric mean. **, p < 0.01; ***, p < 0.001; ****, p < 0.0001. See also Figure S5.

### Metabolism of *Bacteroides*-produced propionate supports *S.* Tm growth *in vivo*

To determine whether *Bacteroides*-produced propionate promotes *S*. Tm intestinal colonization *in vivo*, we investigated the role of *S*. Tm propionate utilization during intestinal infection in a germ-free mouse model. Germ-free mice were left germ-free or were colonized with wildtype *B. theta* or *B. theta* BT1686-89 (**Figure 6A**). In addition, a subset of mice colonized with *B. theta* BT1686-89 were treated with propionate in their drinking water (**Figure 6A**). Measurement of propionate in the feces of mice seven days after colonization revealed that mice given wildtype *B. theta* had significantly higher levels of propionate than mice colonized with *B. theta* BT1686-89 (**Figure 6B**). Supplementation with propionate in the drinking water of mice colonized with *B. theta* BT1686-89 increased the amount of propionate in the feces (**Figure 6B**). Colonization of the three groups with their respective strain of *Bacteroides* was confirmed after seven days (**Figure 6C**). Next, we infected each group with an equal mixture of wildtype *S.* Tm and Δ*prpC*. Wildtype *S.* Tm was able to outcompete the Δ*prpC* mutant in mice colonized with wildtype *B. theta* three days after infection, (**Figure 6D-E**). The wildtype *S*. Tm competitive advantage was abrogated in mice colonized with *B. theta* BT1686-89 but restored if *B. theta* BT1686-89 mice are given propionate in the drinking water (**Figure 6D-E**). All experimental groups had equal levels of intestinal inflammation, revealing that differences in propionate-dependent intestinal colonization by *S*. Tm in mice colonized with wildtype *B. theta* or *B. theta* BT1686-89 were not due to altered host-immune responses (**Figure 6F**). Collectively, these experiments show that *S*. Tm can overcome the inhibitory effects of *Bacteroides*-derived propionate in the inflamed gut.

**Figure 6.**
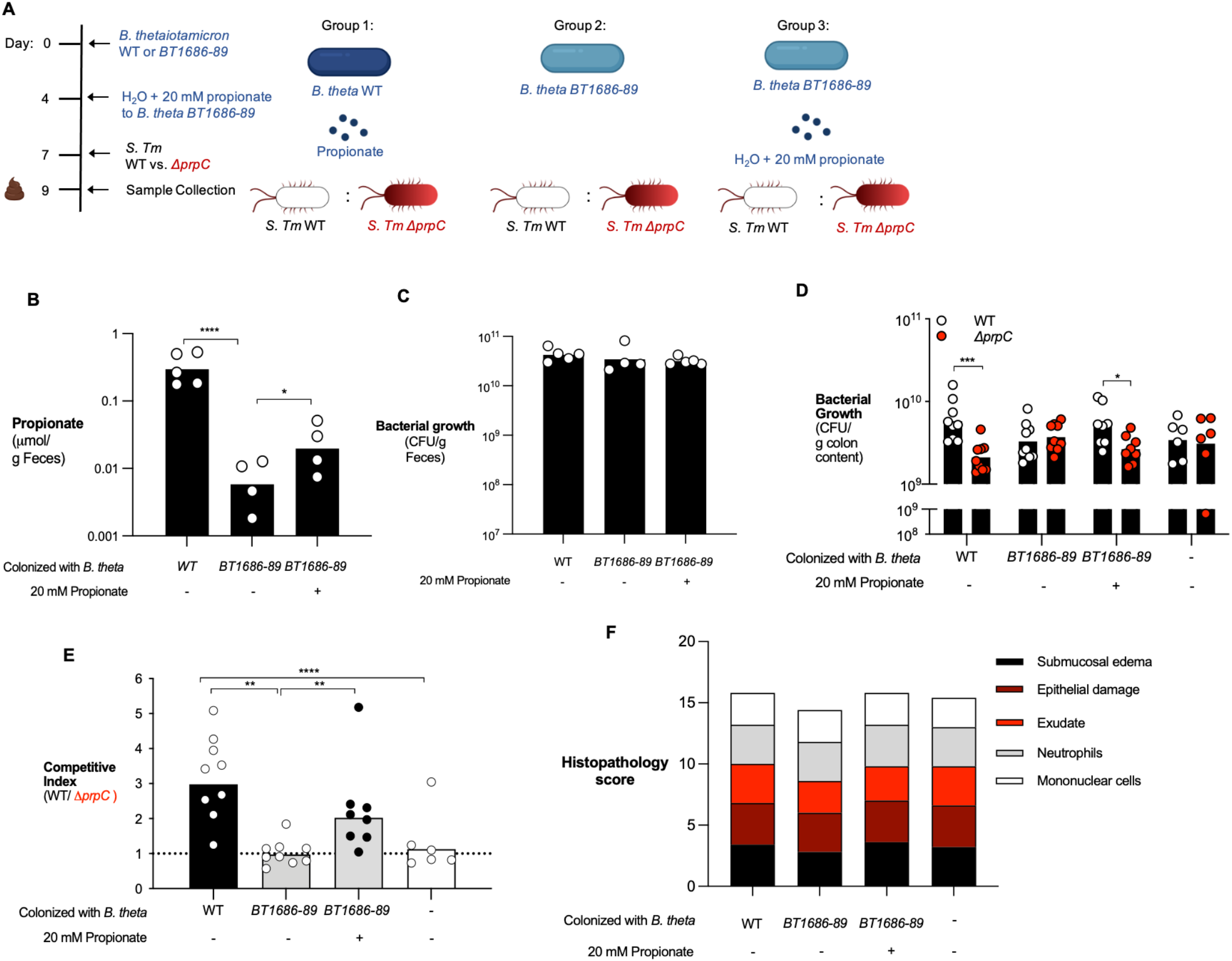
*S.* Tm utilization of microbiota-derived propionate provides an advantage during infection. (A) Schematic representation of the experiment and groups used. Germ-free mice were colonized with either wildtype *B. theta* (WT) or *B. theta* BT1686-89 for 7 days before infection with an equal mixture of wildtype *S.* Tm and a Δ*prpC* mutant. (B) Propionate concentration in the feces was measured after 7 days of colonization with different strains of *Bacteroides* and propionate supplementation. (C) Before infection, fecal samples were collected from monocolonized mice and plated on blood agar to confirm equal colonization between different Bacteroides strains. (D) Monocolonized mice were infected with an equal mixture of the wildtype *S*. Tm and Δ*prpC* mutant. The abundance of each strain in the colon content was determined by selective plating three days post-infection. (E) Competitive index was calculated from CFU counts in (D).(F) Combined histopathology score of pathological lesions in the cecum of mice from (D). Each dot represents one animal. Bars represent the geometric mean. *, p < 0.05; **, p < 0.01; ***, p < 0.001; ****, p < 0.0001.

## Discussion

For decades, the antimicrobial properties of propionate have been leveraged to limit *Salmonellae* infection in agricultural animals (32, 39). However, in this study, we report findings that *S*. Tm can overcome the inhibitory effects of propionate if inflammation-derived nitrate is available. Our data show that *S.* Tm upregulates machinery used to perform propionate catabolism if nitrate is present. This leads to an advantage *in vivo* as strains of *S.* Tm that can perform propionate metabolism have an advantage over strains that cannot catabolize propionate. Collectively, we describe a novel mechanism by which *S.* Tm contends with a component of colonization resistance by performing anaerobic respiration in the inflamed gut (**Figure S4**).

A recent report determined that propionate production by gut microbiota members inhibited *S.* Tm colonization *in vivo* (13). For the study, the researchers utilized a mouse model of chronic *S.* Tm infection, characterized by lower bacterial burden and mild intestinal inflammation (13). In contrast, we investigated the interaction between propionate and *S.* Tm in a mouse model of *S.* Tm-induced gastroenteritis that results in severe inflammation (40) (**Figure 3C-D**). *S.* Tm causes inflammation by invading the intestinal epithelium and triggering an immune response (17). Consequently, innate immune cells release reactive oxygen and nitrogen species that react to generate alternative electron acceptors, including tetrathionate and nitrate (23, 31, 41). The presence of alternative electron acceptors provides *S.* Tm with the opportunity to perform anaerobic respiration and outgrow the obligate anaerobic microbiota (22, 23, 41). Furthermore, the inflamed gut is a unique niche in which *S.* Tm alters its metabolism and begins to utilize carbon sources that require respiration (28). Some of these carbon sources have been identified, including 1,2-propanediol, ethanolamine, and fructose-asparagine (24, 25, 42). However, many remain unknown (28). Here, we show that *S.* Tm can metabolize propionate via anaerobic respiration, and this provides *S.* Tm with an advantage *in vivo.* Interestingly, propionate utilization was specific to the availability of nitrate *in vitro* and *in vivo*, supporting the idea that nitrate is involved in the regulation of the *prpBCDE* operon. By utilizing a different model of *S.* Tm infection, we show a new mechanism by which *S.* Tm mitigates the effects of propionate.

Multiple mechanisms have been proposed by which propionate inhibits *Salmonella* and other enteric pathogens. Initial studies hypothesized that propionate reduced *S.* Tm growth through the generation and accumulation of toxic intermediates (14). Subsequent research focused on the ability of propionate to diffuse into the cytoplasm of the bacteria and decrease intracellular pH (13, 16). This is predicted to impede the ability of *S.* Tm to colonize its host (13). Additional modes of inhibition include repression of *S.* Tm invasion through destabilization of *hilD*, a regulator of *Salmonella* Pathogenicity Island 1 (SPI1) (43). However, past studies that examined the inhibitory effects of propionate did not consider the presence of inflammation-derived alternative electron acceptors. Indeed, we showed that the addition of nitrate to cultures containing propionate increased the growth rate of *S.* Tm despite acidic conditions. Therefore, we propose that inflammation-derived nitrate provides *S.* Tm with the ability to overcome propionate-induced toxicity. The impact of inflammation-dependent propionate catabolism in regulation of *S.* Tm invasion remains to be explored.

By leveraging a mono-colonized germ-free mice model, we determined that the advantage observed for wildtype *S.* Tm over a *prpC* deficient mutant is specific to microbiota-derived propionate. The 2-methylcitrate cycle is fueled by 1,2-propanediol and propionate; both metabolites are produced by members of the microbiota (25). A previous study identified that 1,2-propanediol fuels *S*. Tm growth during infection (25). In this work, we showed that the advantage observed for wildtype *S.* Tm over a *prpC* deficient mutant is specific to propionate and not 1,2-propanediol using germ-free mice colonized with the specific strains of *B. theta.* Future work should examine how *S*. Tm integrates microbiota-derived 1,2-propanediol and propionate to grow *in vivo.*

In conclusion, this study shows that intestinal inflammation enables *S.* Tm to overcome propionate-mediated colonization resistance. Together, our findings provide the paradigm-shifting perspective that, during infection, microbiota-derived propionate may aid in intestinal pathogen colonization. Therefore, in addition to the role of propionate in promoting colonization resistance, we propose that during infection, microbiota-derived propionate may also support *S.* Tm expansion in the intestinal tract. The inhibitory and beneficial effects of propionate during pathogen colonization may have significant implications on the use of microbiota-derived propionate as an antimicrobial treatment during *S.* Tm gastroenteritis and will be an important area of future research.

## Acknowledgments

We would like to thank Dr. Eric Martens for kindly providing *Bacteroides thetaiotaomicron* and *Bacteroides thetaiotaomicron* BT1686-89. C.D.S. was supported by Dorothy Beryl and Theodore Roe Austin Pathology Research Fund and T32AI112541. W.Y. was supported by the Basic Science Research Program through the National Research Foundation of Korea (N.R.F.) by the Ministry of Education 2020R1A6A3A03037326. N.G.S. was supported by T32ES007028-46. N.J.F. was supported by T32DK007673. Work in S.R.’s laboratory was supported by a grant (19162MFDS037) from the Ministry of Food and Drug Safety of Korea, Korea in 2021. Work in M.X.B.’s laboratory was supported by V Scholar grant V2020-013 from The V Foundation for Cancer Research, Vanderbilt Digestive Disease Pilot and Feasibility grant P30 058404, A.C.S. Institutional Research Grant IRG-19-139-59, VICC GI SPORE grant P50CA236733, United States-Israel Binational Science Foundation grant 2019136 and Vanderbilt Institute for Clinical and Translational Research Grant VR53102.

## Author contributions

M.X.B., C.D.S., W.Y., and S.R. designed and conceived the study. C.D.S., W.Y., N. S., T.T., J.Z., N.J.F., D.K., and J.K. performed all experiments. M.W.C. performed propionate measurements. All authors contributed to the data analysis and preparation of the manuscript.

## Declaration of interests

The authors declare no conflict of interest.

## Material and Methods

### Contact for reagent and resource sharing

Further information and requests for resources and reagents should be directed to and will be fulfilled by the Lead Contact, Mariana X. Byndloss. All unique reagents generated in this study are available from the lead contact without restriction.

### Experimental models

#### Mouse husbandry

All animal experiments were approved by the Institution of Animal Care and Use Committee at Vanderbilt University Medical Center. Female C57BL/6J mice and *Nos2*-deficient (on the C57BL/6J background), aged 6 weeks, were obtained from The Jackson Laboratory. Mice were housed in individually ventilated cages with *ad libitum* access to chow and water. Germ-free (G.F.) Swiss Webster mice were initially purchased from Taconic Farms and maintained by the investigators at Vanderbilt University Medical Center. Experiments in this study were performed with 6-week-old male and female G.F. mice.

Animals were randomly assigned to treatment groups before experimentation. At the end of the experiment, mice were humanely euthanized using carbon dioxide inhalation. Animals that had to be euthanized for humane reasons before reaching the predetermined time point were excluded from the analysis.

#### Bacterial culture

*S.* Tm strains (**Table S1**) were routinely grown aerobically at 37 °C in L.B. Broth (10 g/L tryptone, 5 g/L yeast extract, 10 g/L sodium chloride) or on L.B. agar plates (10 g/L tryptone, 5 g/L yeast extract, 10 g/L sodium chloride, 15 g/L agar). When appropriate, L.B. agar plates and broth were supplemented with 100 μg/mL streptomycin (Strep), 30 μg/mL chloramphenicol (Cm), 100 μg/mL carbenicillin (Carb), 50 μg/mL kanamycin (Kan). *B. thetaiotaomicron* strains were cultured in an anaerobic chamber (85% nitrogen, 10% hydrogen, 5% carbon dioxide, Coy Lab Products). *B. thetaiotaomicron* strains (**Table S1**) were routinely cultured on blood agar plates (37 g/L brain heart infusion medium, 15 g/L agar, 50 mL sheep blood).

### Experimental procedure

#### Construction of bacterial strains and plasmids

All *S.* Tm mutants were generated from the SL1344 parent strain. To construct a Δ*prpC* mutant, upstream and downstream regions of approximately 0.5 kb in length were amplified by PCR and then purified. The pRDH10 suicide vector was digested with SalI, purified, and assembled with the *prpC* PCR fragments to form pRDH10∷Δ*prpC*. pRDH10∷Δ*prpC* was then transformed into *E. coli* S17-1 λ*pir*. Conjugation was then performed at 30°C, and exconjugants in which the suicide plasmid had integrated into the chromosome of *S.* Tm were recovered on L.B. agar plates containing streptomycin and chloramphenicol. Subsequent sucrose selection was performed on sucrose plates (5% sucrose, 10 g/L tryptone, 5 g/L yeast extract, 10 g/L sodium chloride, 15 g/l agar) to select for a second crossover events. PCR was performed to detect events that lead to the unmarked deletion of *prpC.* The Δ*prpBCDE* mutant strain was constructed as described above, but with primers designed for regions upstream of *prpB* and downstream of *prpE* in order to create pRDH10∷Δ*prpBCDE*. To generate Δ*invA* Δ*spiB*, *invA* was deleted from SL1344 using the lambda red recombination method (44). The Kan^R^ cassette was amplified from pKD13 using the primers with homology to upstream and downstream of *invA.* The resulting PCR product was integrated into the *invA* region in a wildtype strain containing the plasmid pKD46, followed by the selection of Δ*invA*∷Kan mutants. The Kan^R^ cassette was removed using the plasmid PCP20 (44). The double mutant Δ*invA* Δ*spiB* was then constructed by deleting *spiB* from Δ*invA* using lambda red recombination as detailed previously. To generate Δ*invA* Δ*spiB* Δ*prpC*, *prpC* was deleted from Δ*invA* Δ*spiB* using the lambda red recombination method as described above. To construct a Δ*napA* Δ*narG* Δ*narZ* Δ*prpC* mutant, conjugation was performed with Δ*napA* Δ*narG* Δ*narZ* (22) and S17-1 λ*pir* transformed with pRDH10∷Δ*prpC*. Exconjugants were recovered on L.B. agar plates containing streptomycin, carbenicillin, and chloramphenicol, and then sucrose selection was performed for second crossover events. PCR was performed to detect events that lead to the unmarked deletion of *prpC.* To introduce selectable marker, the *phoN*∷Kan^R^ or *phoN*∷Cm^R^ mutation was transduced by phage P22 HT *int-105* (<u>45</u>) into strains as indicated in Table S1. To complement the *prpC* deletion, the *prpC* gene was amplified and then combined with the BamHI-digested pUHE21-2*lacI*^q^ plasmid using Gibson Assembly. The resulting plasmid was transformed into *E. coli* DH5α, and selected for on L.B. agar plates containing carbenicillin. Plasmid ligation was confirmed by PCR, and pUHE21-2*lacI*^q^∷*prpC* was then transformed into Δ*prpC. B. thetaiotaomicron* containing a deletion in genes BT1686-89 (*B. theta* BT1686-89) was constructed previously (38), and both wildtype *B. thetaiotaomicron* VPI-5482 Δ*tdk* (*B. theta*) (46) and *B. theta* BT1686-89 strains were provided to the investigators by Dr. Eric Martens.

#### *In vitro* growth assays

Non-Carbon E Salts (NCE media) containing 3.94 g/l monopotassium phosphate, 5.9 g/l dipotassium phosphate, 4.68 g/l ammonium sodium hydrogen phosphate tetrahydrate, 2.46 g/l magnesium sulfate heptahydrate, 1 mM magnesium sulfate, 0.1% casamino acids, 1% vitamin and mineral supplements (A.T.C.C.) was supplemented with 40 mM of an electron acceptor (dimethyl sulfoxide (DMSO), trimethylamine N-oxide (TMAO), fumarate, potassium tetrathionate, or sodium nitrate) or a combination of 10 mM propionate and an electron acceptor. Media was placed in the anaerobic chamber 48 hours prior to inoculation. Overnight aerobic cultures of *S.* Tm strains were harvested, washed in PBS, and resuspended in NCE media. wildtype *S.* Tm was then added to anaerobic media containing different electron acceptors (plus or minus propionate) at a final concentration of 1 x 10^4^ CFU/mL. Growth was determined after 24 hours by spreading serial ten-fold dilutions on LB agar plates.

To measure anaerobic growth of wildtype *S.* Tm, Δ*prpC*, and Δ*prpBCDE* with propionate and/or nitrate, overnight cultures of each strain were diluted into anaerobic NCE media containing 40 mM glycerol and sodium nitrate. Strains were incubated for four hours, harvested, washed in PBS, and resuspended in NCE media. In a 96 well plate, strains were added to NCE media containing propionate, nitrate, or a combination of both at a final OD_600_ = 0.001. OD_600_ was measured after 24 hours using the Epoch 2 plate reader (BioTek). Similarly, growth of wildtype *S.* Tm, Δ*prpC*, and Δ*prpBCDE* in NCE media containing 40 mM glycerol and nitrate or 5 mM glucose was determined as described above with the exception that no anaerobic back-dilution was done.

To confirm growth of a complemented Δ*prpC* strain, overnight cultures of wildtype *S.* Tm + pUHE21-2*lacI*^q^, Δ*prpC* + pUHE21-2*lacI*^q^, and Δ*prpC* + pUHE21-2*lacI*^q^∷*prpC* were harvested, washed in PBS, and resuspended in NCE media. Strains were then added to NCE media containing 200 μM isopropyl β-D-1-thiogalactopyranoside (IPTG) and 10 mM propionate or 200 μM IPTG, 10 mM propionate, and 40 mM nitrate at a final O.D. = 0.001. OD_600_ was measured after 24 hours using the Epoch 2 plate reader (BioTek). *In vitro* growth assays were performed in triplicate with different colonies.

#### *In vitro* gene expression

Overnight cultures of wildtype *S.* Tm were harvested and 1 x 10^9^ CFU was added to 5 mL of NCE media or NCE media supplemented with either 10 mM propionate, 40 mM nitrate, or a combination of both. Cultures were incubated for 4 hours prior to R.N.A. extraction (performed according to instructions for SurePrep TrueTotal RNA Purification Kit). R.N.A. (500 ng) was reverse transcribed using an iScript gDNA Clear cDNA synthesis kit (Bio-Rad). Quantitative PCR was performed using S.Y.B.R. green (SsoAdvanced; Bio-Rad) for *prpR, prpB,* and *prpC*. The expression of target genes was normalized to that of *gyrB*. Primers are listed in **Table S2**.

#### Growth and generation time of *S.* Tm at decreasing pH

NCE media was adjusted to pH 7.0, pH 6.5, and pH 6.0 using 10 M hydrochloric acid (HCl). 10 mL of media from each pH was supplemented with 10 mM propionate or 10 mM propionate + 40 mM nitrate and placed in the anaerobic chamber. Overnight cultures of wildtype *S.* Tm and Δ*prpC* were washed in PBS and resuspended in NCE media. Cultures were adjusted to an O.D. = 0.1 and then diluted 1:100 into pH-adjusted media. Samples were incubated at 37 °C, and aliquots were removed every two hours for 14 hours. Aliquots were then serially diluted and plated onto LB agar to determine bacterial numbers. Generation time (G) was calculated according to (47) and using the following formula: G = (T_2_ – T_1_)/(3.3log(B_2_/B_1_) where T_2_ equals the time at the end of exponential phase, and T_1_ is the time at the beginning of the exponential phase. B_2_ corresponds to the bacterial number at T_2,_ and B_1_ equals the bacterial number at T_1_.

#### Growth of *Bacteroides thetaiotaomicron* strains in mucin broth

Porcine mucin was dissolved in 1x NCE salts at a final concentration of 0.5% (w/v). Mucin broth was inoculated with a fresh colony of *B. theta* BT1686-89.and incubated under anaerobic conditions for 96 hours at 37°C or inoculated with wildtype *B. theta* and incubated for 48 hours at 37°C. To measure the growth of *B. theta* BT1686-89, aliquots were removed from cultures, and ten-fold serial dilutions were plated on blood agar plates to calculate bacterial numbers. Digested mucin broth from each strain was filter-sterilized (0.5 μm pore size) before propionate measurements or competitive growth assays.

#### Competitive growth assays

Competition assays were performed in either digested mucin broth or NCE Media. Propionate (10 mM) or nitrate (40 mM) were added as indicated. A 1:1 ratio of two overnight bacterial strains at a final concentration of 1 x 10^4^ CFU/mL were added to the media and incubated anaerobically for 18 hours. Bacterial numbers were determined by plating serial dilutions on selective LB Agar plates. *In vitro* competition assays were performed in triplicate with different colonies

#### Animal experiments

##### Streptomycin-treated mouse model

Groups of 6 – 7-week-old C57BL/6 mice were treated with 5 g/L streptomycin in the drinking water for 48 hours which was then removed 24 hours before infection with *S.* Tm. For competitive infections, mice were orally inoculated with a 1:1 mixture of 1 x 10^9^ CFU of each strain. Fecal samples were collected two days after infection for propionate measurement. Four days after infection, samples for histopathology, cecal tissue for RNA extraction, colonic luminal content for bacterial plating, and cecal content for propionate measurements were collected. In some experiments, mice were given 1 g/L aminoguanidine hydrochloride in their drinking water immediately after infection with *S.* Tm.

##### Germ-Free Swiss Webster mice

Groups of 6-week-old germ-free mice were colonized with approximately 1 x 10^8^ CFU of wildtype *B. theta* or *B. theta* BT1686-89. Four days after colonization a subset of mice colonized with *B. theta* BT1686-89were given 20 mM propionate in the drinking water. After 7 days, fecal samples were collected from mono-colonized mice to measure *B. theta* colonization and propionate levels. Mono-colonized and germ-free mice were then infected with an equal mixture of 1 x 10^7^ CFU of wildtype *S.* Tm and Δ*prpC.* Three days after infection with *S.* Tm samples were collected as described above.

#### Nitrate measurements

Intestinal nitrate measurements were performed as described previously (48). Briefly, mice were euthanized, and the intestine was removed and divided along its sagittal plane. The mucus layer was gently scraped from the tissue and homogenized in 200 μl PBS and then placed on ice. Samples were centrifuged at 5,000 × g for 10 min at 4°C to remove the remaining solid particles. The supernatant was then filter sterilized (0.2-μm pore size). Measurement of intestinal nitrate followed an adaptation of the Griess assay. In this assay, nitrate was first reduced to nitrite by combining 50 μl of each sample with 50 μl of Griess reagent 1 containing vanadium(III) chloride (0.5 M HCl, 0.2 mM VCl3, 1% sulfanilamide), and then the mixture was incubated at room temperature for 10 min. Next, 50 μl of Griess reagent 2 [0.1% (1-naphthyl)ethylenediamine dichloride] was added to each sample. Absorbance at 540 nm was measured immediately after the addition of Griess reagent 2 to detect any nitrite present in the samples. The samples were then incubated for 8 h at room temperature (to allow for reduction of nitrate to nitrite), and the absorbance at 540 nm was measured again. The initial absorbance (prior to reducing nitrate to nitrite) was subtracted from the absorbance after 8 h to determine nitrate concentrations in the mucus layer.

#### Quantification of *Nos2* expression by qRT-PCR

Cecal tissue was homogenized using a FastPrep-24 and RNA extracted using the TRI reagent method. RNA (1 μg) was reverse transcribed using an iScript gDNA Clear cDNA synthesis kit (Bio-Rad). Quantitative PCR was performed with SYBR green (SsoAdvanced; Bio-Rad) for *Nos2* (Primers listed in **Table S2**). The expression of *Nos2* was normalized to the housekeeping gene *Act2b*, encoding β-actin.

#### Propionate measurements

##### Extraction and normalization

Fecal matter was weighed, diluted to a final density of 0.125 g/mL in MeOH/H_2_O (1:5) and homogenized with a cordless *Pellet Pestle* tissue grinder equipped with disposable polypropylene mixers (Fisher). Insoluble debris was removed by centrifugation (12,000 x g, 30 min, 5 °C); the supernatants were transferred to clean Eppendorf tubes and stored at −20 °C until the day of analysis.

##### Propionate Analysis

Propionate was derivatized with the reagent dansylhydrazine and the carboxyl activating agent 1-Ethyl-3-(3-dimethylaminopropyl)carbodiimide (EDC) and measured as its corresponding dansylhydrazone derivative (49). Briefly, fecal extracts (10 μL) were spiked with a stable isotope-labeled internal standard propionate-d_5_ (1 nmol) and derivatized in H_2_O/DMSO (2:1) containing 50 mM sodium phosphate buffer (pH = 4), 12.5 mg/mL dansylhydrazine, and 12.5 mg/mL EDC. Due to the limited stability of EDC in water, stock solutions of EDC should be made up in ice-cold water and used immediately. After two hours at room temperature, dansylated derivatives were extracted once with ethyl acetate (750 μL). The organic (top) layer was transferred to a clean Eppendorf tube, dried under a gentle stream of nitrogen gas, and reconstituted in 150 μL of acetonitrile/water (1:1) prior to analysis. Calibration standards for unlabeled propionate were prepared in water, derivatized, and extracted in the same manner. LC-MS/MS analysis was performed using a Thermo TSQ Quantum mass spectrometer interfaced to a Thermo HTC PAL refrigerated autosampler and a Thermo Surveyor HPLC pump. A Waters XTerra MS analytical column (2.1 mm x 100 mm, 3.5 μm) was used for all chromatographic separations. Mobile phases were made up of 0.2% acetic acid and 15 mM ammonium acetate in (A) H_2_O/CH_3_CN (9:1) and in (B) CH_3_CN/CH_3_OH/H_2_O (90:5:5). Gradient conditions were as follows: 0–1 min, B = 0 %; 1–8 min, B = 0–100 %; 8–10 min, B = 100 %; 10–10.5 min, B = 100–0 %; 10.5–15 min, B = 0 %. The flow rate was maintained at 300 μL/min; a software-controlled divert valve was used to transfer eluent from 0–2.0 min of each chromatographic run to waste. The total chromatographic run time was 15 min. The autosampler tray temperature and the column compartment temperature were maintained at 5 °C and 50 °C respectively. The sample injection volume was 10 μL. The autosampler injection valve and the sample injection needle were flushed and washed sequentially with mobile phase B (two cycles) and mobile phase A (two cycles) between each injection. The mass spectrometer was operated in positive ion mode. Quantitation was based on single reaction monitoring detection of the following dansylated analogues: propionate, *m/z* 322 → 235, C.E. 15; propionate-d_5_: *m/z* 327 → 235, C.E. 15. The following optimized source parameters were used for the detection of analytes and internal standards. N_2_ sheath gas 40 psi; N_2_ auxiliary gas 5 psi; spray voltage 4 kV; capillary temperature 300 °C; tube lens voltage 120 V; declustering voltage 20 V. Data acquisition and quantitative spectral analysis were done using Thermo-Finnigan Xcalibur version 2.0.7 SP1 and Thermo-Finnigan LCQuan version 2.7, respectively. Calibration curves were constructed by plotting peak area ratios (analyte / internal standard) against analyte concentrations for a series of nine calibration standards, ranging from 0.01 to 100 nmol propionate. A weighting factor of 1/C^2^ was applied in the linear least-squares regression analysis to maintain homogeneity of variance across the concentration range (%RE ≤ 15% at C > LLOQ).

#### Histopathology scoring

Formalin fixed cecal tissue sections were stained with hematoxylin and eosin, and a veterinary pathologist performed a blinded evaluation using criteria shown in **Table S3** as described previously (50). Representative images were taken using a Leica DM750 microscope and a Leica ICC50W camera.

#### Quantification and statistical analysis

Statistical data analysis was performed using Graphpad PRISM. Fold changes of ratios (bacterial competitive index and mRNA levels), and bacterial numbers were transformed logarithmically prior to statistical analysis. An unpaired Student’s t test was used on the transformed data to determine whether differences between groups were statistically significant (p < 0.05). When more than two treatments were used, statistically significant differences between groups were determined by one-way ANOVA followed by Tukey’s HSD test (between > 2 groups). Significance of differences in histopathology was determined by a one-tailed non-parametric test (Mann-Whitney).

## SUPPLEMENTARY MATERIAL

**Supplemental Table 1.** Bacterial strains were used in this study.

| Bacterial strains | Source |
| --- | --- |
| *S. Typhimurium*, SL1344 Strep<sup>R</sup> | 42 |
| *S. Typhimurium*, $\Delta prpC$ | This study |
| *S. Typhimurium*, $\Delta prpC$ *phoN*::Kan<sup>R</sup> | This study |
| *S. Typhimurium*, $\Delta prpBCDE$ | This study |
| *S. Typhimurium*, $\Delta prpBCDE$ *phoN*::Kan<sup>R</sup> | This study |
| *S. Typhimurium*, $\Delta invA$ | This study |
| *S. Typhimurium*, $\Delta invA \Delta spiB$ | This study |
| *S. Typhimurium*, $\Delta invA \Delta spiB \Delta prpC$ | This study |
| *S. Typhimurium*, $\Delta invA \Delta spiB \Delta prpC$ *phoN*::Kan<sup>R</sup> | This study |
| *S. Typhimurium*, $\Delta napA \Delta narZ$ *narG*::pCAL | 22 |
| *S. Typhimurium*, $\Delta napA \Delta narZ$ *narG*::pCAL5 *phoN*::Kan<sup>R</sup> | 22 |
| *S. Typhimurium*, $\Delta napA \Delta narZ$ *narG*::pCAL5 $\Delta prpC$ | This study |
| *S. Typhimurium*, $\Delta napA \Delta narZ$ *narG*::pCAL5 $\Delta prpC$ *phoN*::Cm<sup>R</sup> | This study |
| *E. coli*, S17-1 $\lambda pir$, *zxx*::RP4 2-(TetR::Mu) (KanR::Tn7) $\lambda pir$ *recA1 thi pro hsdR* (r-m+) | 45 |
| *B. thetaiotaomicron*, VPI 5482 $\Delta tdk$ | 46 |
| *B. thetaiotaomicron*, $\Delta tdk$ $\Delta 1686-1689$ | 38 |

**Supplemental Table 2.**
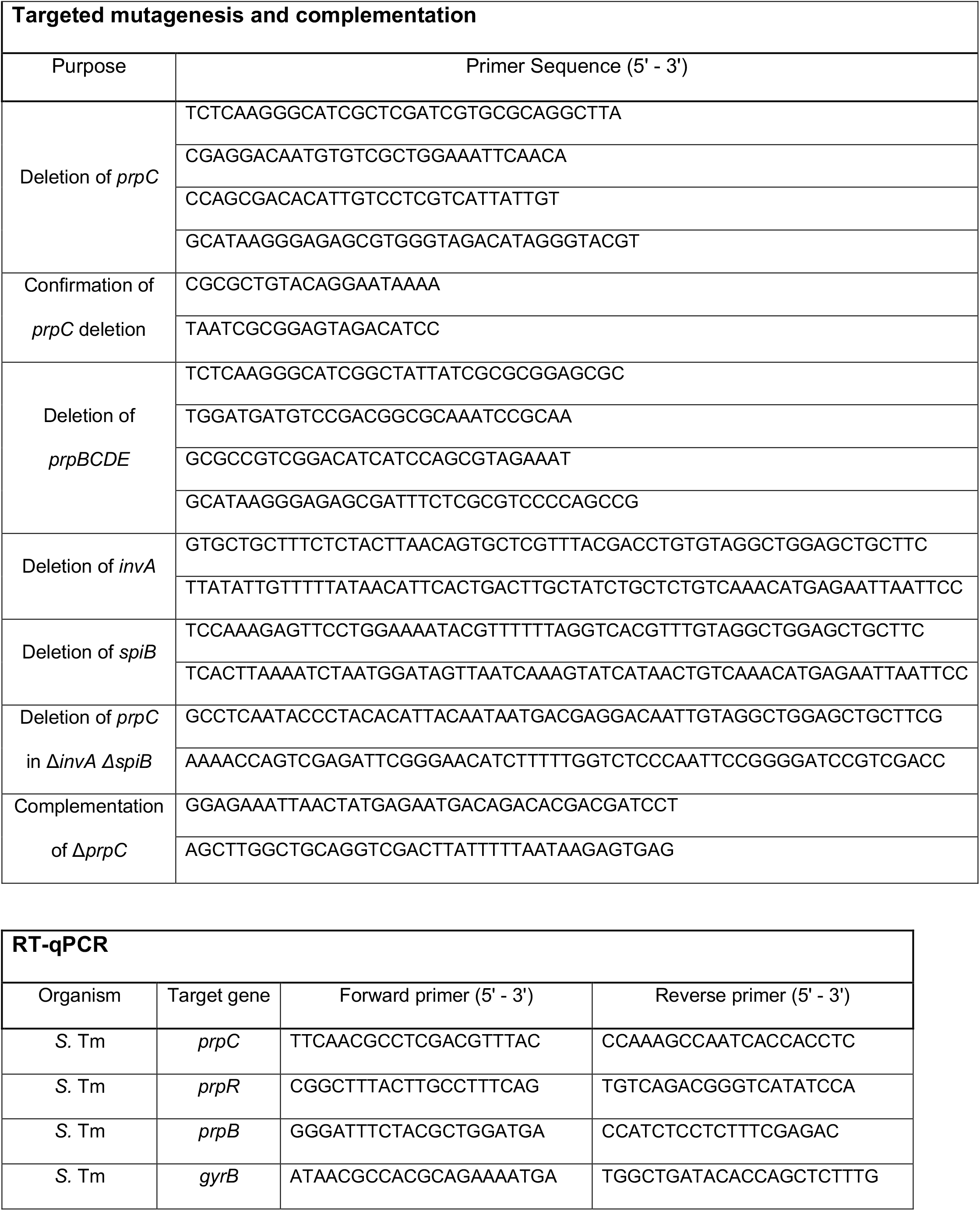
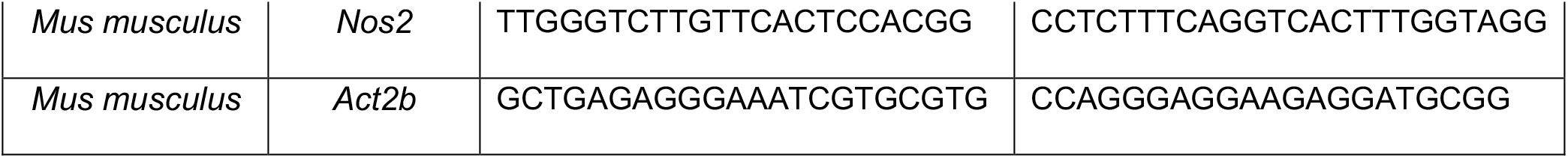
Primers used in this study.

**Supplemental Table 3.** Criteria for blinded scoring of histopathological changes.

| **Score** | **Submucosal edema** | **Epithelial damage** | **Exudate** | **Submucosal neutrophil infiltration (cells/high power field)** | **Submucosal mononuclear cell infiltration (cells/high power field)** |
| --- | --- | --- | --- | --- | --- |
| 0 | No changes | No changes | No changes | No changes (0-5) | No changes (0-5) |
| 1 | Detectable (<10%) | Desquamation | Slight focal accumulation | (6-20) | (5-10) |
| 2 | Mild (10-20%) | Mild erosion | Mild focal accumulation | (21-60) | (10-20) |
| 3 | Moderate (20-40%) | Marked erosion and/or mild ulceration | Moderate multifocal accumulation | (61-100) | (20-40) |
| 4 | Marked (>40%) | Multifocal ulceration | Marked diffuse accumulation | (>100) | (>40) |

**Figure S1.**
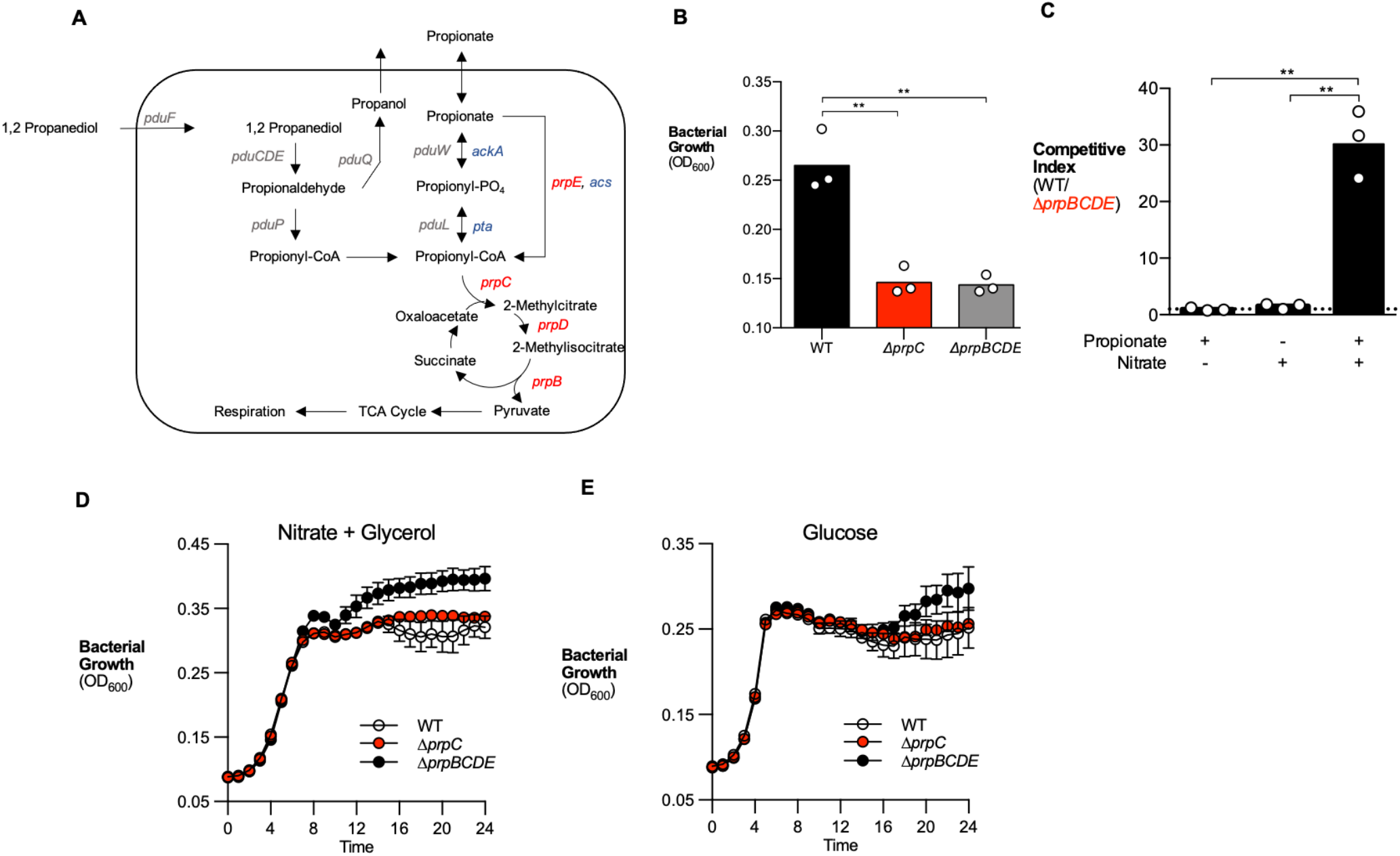
Disruption to *prpBCDE* operon prevents propionate catabolism during anaerobic respiration. (A) Complete model of propionate catabolism in *S.* Tm. Genes in the *prpBCDE* operon (red) metabolize propionate into pyruvate. The intermediate propionyl-CoA is also produced by 1, 2 propanediol catabolism (pdu operon (grey). *acs,* acetyl-CoA synthetase, *ackA*, acetate kinase, *pta,* phosphotransferase (blue). (B) NCE minimal media containing 10 mM propionate and 40 mM nitrate was inoculated with wildtype *S.* Tm, Δ*prpC,* or Δ*prpBCDE*. OD_600_ of each strain was measured after 24 hours of anaerobic growth. (C) NCE minimal media containing 10 mM propionate, 40 mM nitrate, 10 mM propionate and 40 mM nitrate was inoculated with equal mixture of the *S.* Tm wildtype strain and Δ*prpBCDE.* The competitive index was determined after 24 hours of anaerobic growth. (D – E) NCE minimal media containing 40 mM glycerol and 40 mM nitrate (D) or NCE minimal media containing 5 mM glucose (E) were inoculated with wildtype *S.* Tm, Δ*prpC,* or Δ*prpBCDE.* OD_600_ of each strain was measured after 24 hours of anaerobic growth. Each dot represented one biological replicate. Bars represent the geometric mean. **, p < 0.01.

**Figure S2.**
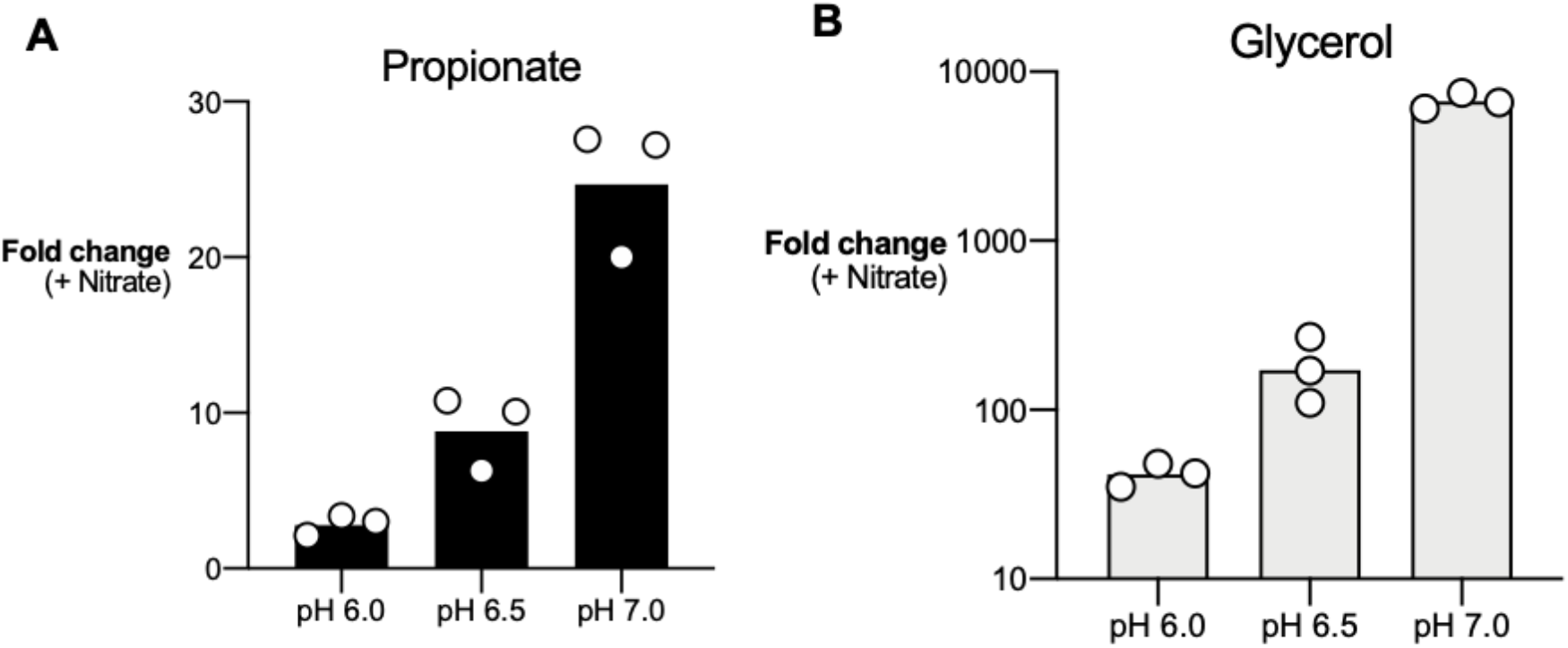
Low pH decreases *S.* Tm growth during anaerobic respiration. (A) NCE minimal media was adjusted to pH 6.0, 6.5, and 7.0 and either 10 mM propionate or 10 mM propionate + 40 mM nitrate was added. Media was inoculated with wildtype *S.* Tm and grown anaerobically for 24 hours. Fold change calculated by comparing growth of *S.* Tm in media containing propionate alone to *S.* Tm grown with both propionate and nitrate. (B) NCE minimal media was adjusted to pH 6.0, 6.5, and 7.0 and either 40 mM glycerol or 40 mM glycerol + 40 mM nitrate was added. Media was inoculated with wildtype *S.* Tm and grown anaerobically for 24 hours. Fold change calculated by comparing growth of *S.* Tm in media containing glycerol alone to *S.* Tm grown with both glycerol and nitrate. Each dot represents one biological replicate. Bars represent the geometric mean.

**Figure S3.**
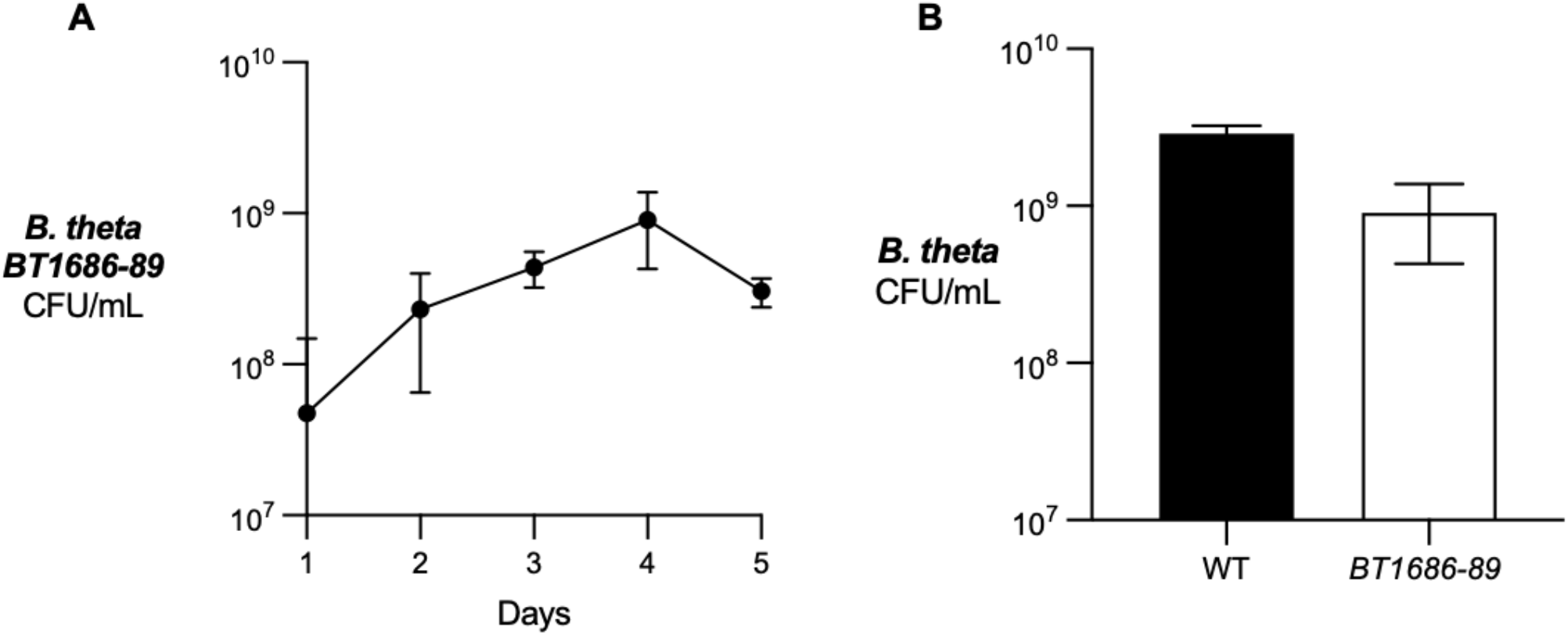
*In vitro* growth of *Bacteroides* strains. (A) Mucin broth was inoculated with *B. thetaiotaomicron* BT1686-89 and aliquots taken every 24 hours for plating on blood agar. Dots represent mean, error bars represent SD (n = 4). (B) Mucin broth was inoculated with wildtype *B. thetaiotaomicron (*WT) *or B. thetaiotaomicron BT161-89* (BT161-89*). B. theta* BT161-89 was cultured anaerobically for 4 days and wildtype *B. theta* was cultured for 2 days. Growth determined by plating on blood agar. Bars represent mean, error bars represent SD (n = 3).

**Figure S4.**
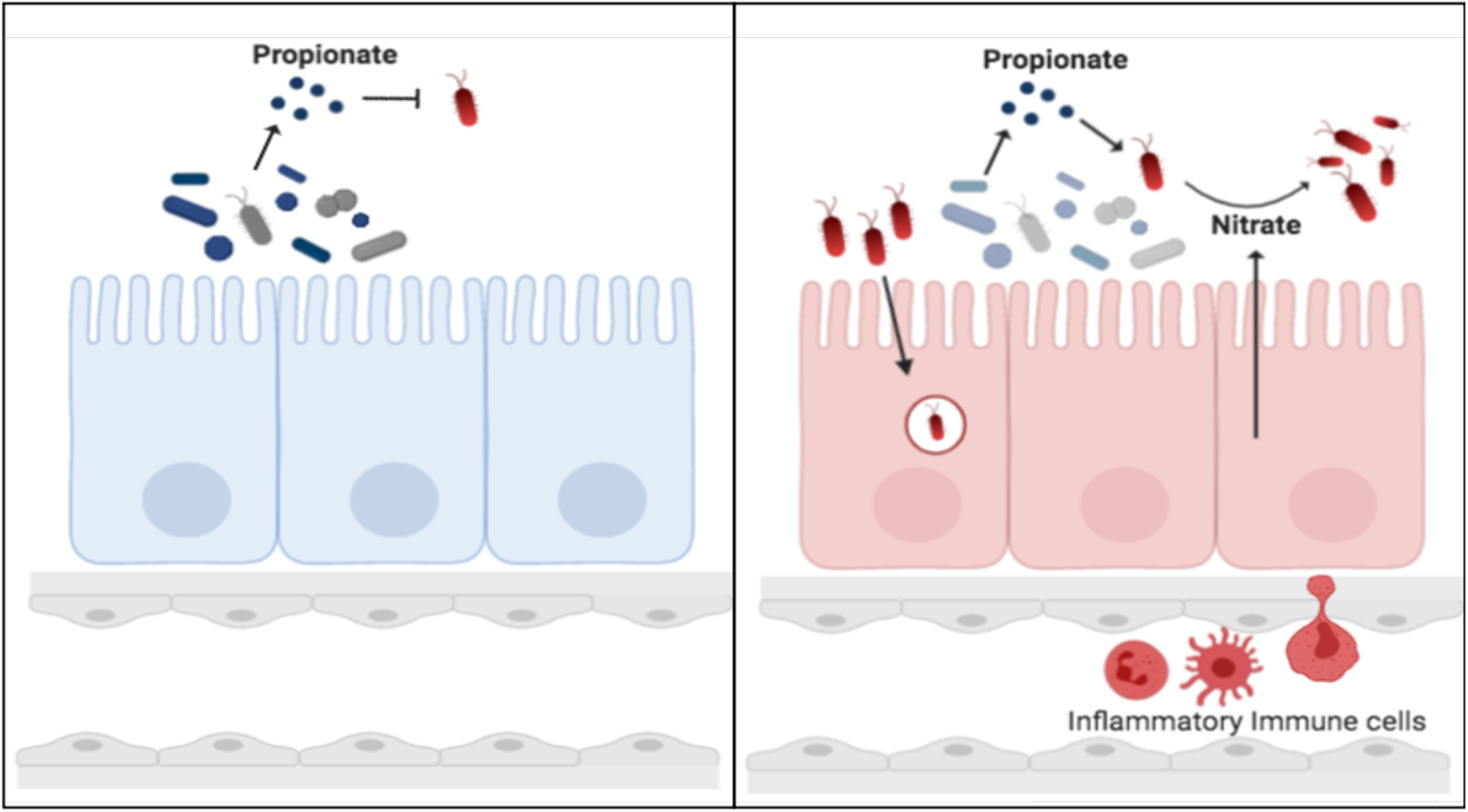
Schematics of S. Tm propionate utilization in the inflamed gut. Upon infection, *S.* Tm (red) uses its’ T3SS-1 to invade the intestinal epithelium. Innate immune cells (red) recognize this invasion and produce nitric oxide (NO) that can be converted into nitrate (NO_3_-). Propionate is produced by microbiota members (blue and gray). We propose *S.* Tm overcomes microbiota-mediated colonization resistance by using propionate as a carbon source to respire via alternative electron acceptors.

## Notes

### Competing Interest Statement

The authors have declared no competing interest.

## REFERENCES

1. Sonnenburg, J.L., Xu, J., Leip, D.D., Chen, C.H., Westover, B.P., Weatherford, J., Buhler, J.D. and Gordon, J.I., 2005. Glycan foraging in vivo by an intestine-adapted bacterial symbiont. Science, 307(5717), pp.1955–1959.

2. Bäckhed, F., Ding, H., Wang, T., Hooper, L. V., Koh, G. Y., Nagy, A., Semenkovich, C. F., & Gordon, J. I. (2004). The gut microbiota as an environmental factor that regulates fat storage. Proceedings of the National Academy of Sciences of the United States of America, 101(44), 15718–15723. https://doi.org/10.1073/pnas.0407076101

3. Lathrop, S. K., Bloom, S. M., Rao, S. M., Nutsch, K., Lio, C. W., Santacruz, N., Peterson, D. A., Stappenbeck, T. S., & Hsieh, C. S. (2011). Peripheral education of the immune system by colonic commensal microbiota. Nature, 478(7368), 250–254. https://doi.org/10.1038/nature10434

4. Zheng, D., Liwinski, T., & Elinav, E. (2020). Interaction between microbiota and immunity in health and disease. Cell research, 30(6), 492–506. https://doi.org/10.1038/s41422-020-0332-7

5. van der Waaij, D., Berghuis-de Vries, J. M., & Lekkerkerk Lekkerkerk-v (1971). Colonization resistance of the digestive tract in conventional and antibiotic-treated mice. The Journal of hygiene, 69(3), 405–411. https://doi.org/10.1017/s0022172400021653

6. Bohnhoff, M., Miller, C. P., & Martin, W. R. (1964). Resistance of the mouse’s intestinal tract to experimental Salmonella Infection. I. Factors which interfere with the initiation of off infection by oral inoculation. The Journal of experimental medicine, 120(5), 805–816. https://doi.org/10.1084/jem.120.5.805

7. Sassone-Corsi, M., & Raffatellu, M. (2015). No Vacancy: How Beneficial Microbes Cooperate with Immunity To Provide Colonization Resistance to Pathogens. The Journal of Immunology. https://doi.org/10.4049/jimmunol.1403169

8. Petersson, J., Schreiber, O., Hansson, G. C., Gendler, S. J., Velcich, A., Lundberg, J. O., Roos, S., Holm, L., & Phillipson, M. (2011). Importance and regulation of the colonic mucus barrier in a mouse model of colitis. American Journal of Physiology - Gastrointestinal and Liver Physiology. https://doi.org/10.1152/ajpgi.00422.2010

9. Ivanov, I. I., Atarashi, K., Manel, N., Brodie, E. L., Shima, T., Karaoz, U., Wei, D., Goldfarb, K. C., Santee, C. A., Lynch, S. V., Tanoue, T., Imaoka, A., Itoh, K., Takeda, K., Umesaki, Y., Honda, K., & Littman, D. R. (2009). Induction of Intestinal Th17 Cells by Segmented Filamentous Bacteria. Cell. https://doi.org/10.1016/j.cell.2009.09.033

10. Benson, A., Pifer, R., Behrendt, C. L., Hooper, L. V., & Yarovinsky, F. (2009). Gut Commensal Bacteria Direct a Protective Immune Response against Toxoplasma gondii. Cell Host and Microbe. https://doi.org/10.1016/j.chom.2009.06.005

11. Deriu, E., Liu, J. Z., Pezeshki, M., Edwards, R. A., Ochoa, R. J., Contreras, H., Libby, S. J., Fang, F. C., & Raffatellu, M. (2013). Probiotic bacteria reduce Salmonella typhimurium intestinal colonization by competing for iron. Cell Host and Microbe. https://doi.org/10.1016/j.chom.2013.06.007

12. Sorbara, M. T., & Pamer, E. G. (2019). Interbacterial mechanisms of colonization resistance and the strategies pathogens use to overcome them. Mucosal immunology, 12(1), 1–9. https://doi.org/10.1038/s41385-018-0053-0

13. Jacobson, A., Lam, L., Rajendram, M., Tamburini, F., Honeycutt, J., Pham, T., Van Treuren, W., Pruss, K., Stabler, S. R., Lugo, K., Bouley, D. M., Vilches-Moure, J. G., Smith, M., Sonnenburg, J. L., Bhatt, A. S., Huang, K. C., & Monack, D. (2018). A Gut Commensal-Produced Metabolite Mediates Colonization Resistance to Salmonella Infection. Cell Host and Microbe. https://doi.org/10.1016/j.chom.2018.07.002

14. Horswill, A. R., Dudding, A. R., & Escalante-Semerena, J. C. (2001). Studies of Propionate Toxicity in Salmonella enterica Identify 2-Methylcitrate as a Potent Inhibitor of Cell Growth. Journal of Biological Chemistry. https://doi.org/10.1074/jbc.M100244200

15. Byrne, B. M., & Dankert, J. (1979). Volatile fatty acids and aerobic flora in the gastrointestinal tract of mice under various conditions. Infection and immunity, 23(3), 559–563. https://doi.org/10.1128/IAI.23.3.559-563.1979

16. den Besten, G., van Eunen, K., Groen, A. K., Venema, K., Reijngoud, D. J., & Bakker, B. M. (2013). The role of short-chain fatty acids in the interplay between diet, gut microbiota, and host energy metabolism. Journal of lipid research, 54(9), 2325–2340. https://doi.org/10.1194/jlr.R036012

17. Gala ́n JE, Curtiss R III. 1989. Cloning and molecular characterization of genes whose products allow *Salmonella typhimurium* to penetrate tissue culture cells. P.N.A.S. 86:6383–87

18. Alam, M. S., Akaike, T., Okamoto, S., Kubota, T., Yoshitake, J., Sawa, T., Miyamoto, Y., Tamura, F., & Maeda, H. (2002). Role of nitric oxide in host defense in murine salmonellosis as a function of its antibacterial and antiapoptotic activities. Infection and immunity, 70(6), 3130–3142. https://doi.org/10.1128/iai.70.6.3130-3142.2002

19. Szabó, C., Ischiropoulos, H., & Radi, R. (2007). Peroxynitrite: biochemistry, pathophysiology and development of therapeutics. Nature reviews. Drug discovery, 6(8), 662–680. https://doi.org/10.1038/nrd2222

20. Rivera-Chávez, F., & Bäumler, A. J. (2015). The Pyromaniac Inside You: Salmonella Metabolism in the Host Gut. Annual review of microbiology, 69, 31–48. https://doi.org/10.1146/annurev-micro-091014-104108

21. Barrett, E. L., & Riggs, D. L. (1982). Evidence of a second nitrate reductase activity that is distinct from the respiratory enzyme in Salmonella typhimurium. Journal of bacteriology, 150(2), 563–571. https://doi.org/10.1128/JB.150.2.563-571.1982

22. Lopez, C. A., Winter, S. E., Rivera-Chávez, F., Xavier, M. N., Poon, V., Nuccio, S. P., Tsolis, R. M., & Bäumler, A. J. (2012). Phage-mediated acquisition of a type III secreted effector protein boosts growth of Salmonella by nitrate respiration. mBio, 3(3), e00143–12. https://doi.org/10.1128/mBio.00143-12

23. McLaughlin, P. A., Bettke, J. A., Tam, J. W., Leeds, J., Bliska, J. B., Butler, B. P., & van der Velden, A. W. M. (2019). Inflammatory monocytes provide a niche for Salmonella expansion in the lumen of the inflamed intestine. PLoS Pathogens. https://doi.org/10.1371/journal.ppat.1007847

24. Thiennimitr, P., Winter, S. E., Winter, M. G., Xavier, M. N., Tolstikov, V., Huseby, D. L., Sterzenbach, T., Tsolis, R. M., Roth, J. R., & Bäumler, A. J. (2011). Intestinal inflammation allows Salmonella to use ethanolamine to compete with the microbiota. Proceedings of the National Academy of Sciences of the United States of America, 108(42), 17480–17485. https://doi.org/10.1073/pnas.1107857108

25. Faber, F., Thiennimitr, P., Spiga, L., Byndloss, M. X., Litvak, Y., Lawhon, S., Andrews-Polymenis, H. L., Winter, S. E., & Bäumler, A. J. (2017). Respiration of Microbiota-Derived 1,2-propanediol Drives Salmonella Expansion during Colitis. PLoS pathogens, 13(1), e1006129. https://doi.org/10.1371/journal.ppat.1006129

26. Horswill, A. R., & Escalante-Semerena, J. C. (1997). Propionate catabolism in Salmonella typhimurium LT2: Two divergently transcribed units comprise the prp locus at 8.5 centisomes, prpR encodes a member of the sigma-54 family of activators, and the prpBCDE genes constitute an operon. Journal of Bacteriology. https://doi.org/10.1128/jb.179.3.928-940.1997

27. Horswill, A. R., & Escalante-Semerena, J. C. (1999). Salmonella typhimurium LT2 catabolizes propionate via the 2-methylcitric acid cycle. Journal of Bacteriology. https://doi.org/10.1128/jb.181.18.5615-5623.1999

28. Nuccio, S. P., & Bäumler, A. J. (2014). Comparative analysis of Salmonella genomes identifies a metabolic network for escalating growth in the inflamed gut. mBio, 5(2), e00929–14. https://doi.org/10.1128/mBio.00929-14

29. Rondon, M. R., Horswill, A. R., & Escalante-Semerena, J. C. (1995). D.N.A. polymerase I function is required for the utilization of ethanolamine, 1,2-propanediol, and propionate by Salmonella typhimurium LT2. Journal of bacteriology, 177(24), 7119–7124. https://doi.org/10.1128/jb.177.24.7119-7124.1995

30. Friedman, E. S., Bittinger, K., Esipova, T. V., Hou, L., Chau, L., Jiang, J., Mesaros, C., Lund, P. J., Liang, X., FitzGerald, G. A., Goulian, M., Lee, D., Garcia, B. A., Blair, I. A., Vinogradov, S. A., & Wu, G. D. (2018). Microbes vs. chemistry in the origin of the anaerobic gut lumen. Proceedings of the National Academy of Sciences of the United States of America, 115(16), 4170–4175. https://doi.org/10.1073/pnas.1718635115

31. Winter, S. E., Winter, M. G., Xavier, M. N., Thiennimitr, P., Poon, V., Keestra, A. M., Laughlin, R. C., Gomez, G., Wu, J., Lawhon, S. D., Popova, I. E., Parikh, S. J., Adams, L. G., Tsolis, R. M., Stewart, V. J., & Bäumler, J. (2013). Host-derived nitrate boosts growth of E. coli in the inflamed gut. Science. https://doi.org/10.1126/science.1232467

32. Ricke S. C. (2003). Perspectives on the use of organic acids and short chain fatty acids as antimicrobials. Poultry science, 82(4), 632–639. https://doi.org/10.1093/ps/82.4.632

33. Sorbara, M. T., Dubin, K., Littmann, E. R., Moody, T. U., Fontana, E., Seok, R., Leiner, I. M., Taur, Y., Peled, J. U., Van Den Brink, M. R. M., Litvak, Y., Bäumler, A. J., Chaubard, J. L., Pickard, A. J., Cross, J. R., & Pamer, E. G. (2019). Inhibiting antibiotic-resistant Enterobacteriaceae by microbiota-mediated intracellular acidification. Journal of Experimental Medicine. https://doi.org/10.1084/jem.20181639

34. Coombes, B. K., Coburn, B. A., Potter, A. A., Gomis, S., Mirakhur, K., Li, Y., & Finlay, B. B. (2005). Analysis of the contribution of Salmonella pathogenicity islands 1 and 2 to enteric disease progression using a novel bovine ileal loop model and a murine model of infectious enterocolitis. Infection and immunity, 73(11), 7161–7169. https://doi.org/10.1128/IAI.73.11.7161-7169.2005

35. Raffatellu, M., George, M. D., Akiyama, Y., Hornsby, M. J., Nuccio, S. P., Paixao, T. A., Butler, B. P., Chu, H., Santos, R. L., Berger, T., Mak, T. W., Tsolis, R. M., Bevins, C. L., Solnick, J. V., Dandekar, S., & Bäumler, J. (2009). Lipocalin-2 resistance confers an advantage to Salmonella enterica serotype Typhimurium for growth and survival in the inflamed intestine. Cell host & microbe, 5(5), 476–486. https://doi.org/10.1016/j.chom.2009.03.011

36. Lopez, C. A., Rivera-Chávez, F., Byndloss, M. X., & Bäumler, A. J. (2015). The Periplasmic Nitrate Reductase NapABC Supports Luminal Growth of Salmonella enterica Serovar Typhimurium during Colitis. Infection and immunity, 83(9), 3470–3478. https://doi.org/10.1128/IAI.00351-15

37. Reichardt, N., Duncan, S. H., Young, P., Belenguer, A., McWilliam Leitch, C., Scott, K. P., Flint, H. J., & Louis, P. (2014). Phylogenetic distribution of three pathways for propionate production within the human gut microbiota. The I.S.M.E. journal, 8(6), 1323–1335. https://doi.org/10.1038/ismej.2014.14

38. Kovatcheva-Datchary, P., Nilsson, A., Akrami, R., Lee, Y. S., De Vadder, F., Arora, T., Hallen, A., Martens, E., Björck, I., & Bäckhed, F. (2015). Dietary Fiber-Induced Improvement in Glucose Metabolism Is Associated with Increased Abundance of Prevotella. Cell metabolism, 22(6), 971–982. https://doi.org/10.1016/j.cmet.2015.10.001

39. Iba, A. M., & Berchieri, A., Jr (1995). Studies on the use of a formic acid-propionic acid mixture (Bio-add) to control experimental Salmonella infection in broiler chickens. Avian pathology: journal of the W.V.P.A., 24(2), 303–311. https://doi.org/10.1080/03079459508419071

40. Barthel, M., Hapfelmeier, S., Quintanilla-Martínez, L., Kremer, M., Rohde, M., Hogardt, M., Pfeffer, K., Rüssmann, H., & Hardt, W. D. (2003). Pretreatment of mice with streptomycin provides a Salmonella enterica serovar Typhimurium colitis model that allows analysis of both pathogen and host. Infection and immunity, 71(5), 2839–2858. https://doi.org/10.1128/iai.71.5.2839-2858.2003

41. Winter, S. E., Thiennimitr, P., Winter, M. G., Butler, B. P., Huseby, D. L., Crawford, R. W., Russell, J. M., Bevins, C. L., Adams, L. G., Tsolis, R. M., Roth, J. R., & Bäumler, A. J. (2010). Gut inflammation provides a respiratory electron acceptor for Salmonella. Nature, 467(7314), 426–429. https://doi.org/10.1038/nature09415

42. Ali, M. M., Newsom, D. L., González, J. F., Sabag-Daigle, A., Stahl, C., Steidley, B., Dubena, J., Dyszel, J. L., Smith, J. N., Dieye, Y., Arsenescu, R., Boyaka, P. N., Krakowka, S., Romeo, T., Behrman, E. J., White, P., & Ahmer, B. M. (2014). Fructose-asparagine is a primary nutrient during growth of Salmonella in the inflamed intestine. PLoS pathogens, 10(6), e1004209. https://doi.org/10.1371/journal.ppat.1004209

43. Hung, C. C., Garner, C. D., Slauch, J. M., Dwyer, Z. W., Lawhon, S. D., Frye, J. G., McClelland, M., Ahmer, B. M., & Altier, C. (2013). The intestinal fatty acid propionate inhibits Salmonella invasion through the post-translational control of HilD. Molecular Microbiology, 87(5), 1045–1060. https://doi.org/10.1111/mmi.12149

44. Datsenko, K. A. & Wanner, B. L. One-step inactivation of chromosomal genes in Escherichia coli K-12 using PCR products. Proc. Natl. Acad. Sci. USA.97, 6640–6645, doi: 10.1073/pnas.120163297 (2000).

45. Schmieger H. Phage P22-mutants with increased or decreased transduction abilities. Mol Gen Genet. 1972;119:75–88

46. Koropatkin, N. M., Martens, E. C., Gordon, J. I., & Smith, T. J. (2008). Starch catabolism by a prominent human gut symbiont is directed by the recognition of amylose helices. Structure (London, England : 1993), 16(7), 1105–1115. https://doi.org/10.1016/j.str.2008.03.017

47. Todar, K. G., & University of Wisconsin--Madison,. (2006). Todar’s Online textbook of bacteriology. Madison, WI: Kenneth Todar, University of Wisconsin-Madison Dept. of Bacteriology.

48. Byndloss, M. X., Olsan, E. E., Rivera-Chávez, F., Tiffany, C. R., Cevallos, S. A., Lokken, K. L., Torres, T. P., Byndloss, A. J., Faber, F., Gao, Y., Litvak, Y., Lopez, C. A., Xu, G., Napoli, E., Giulivi, C., Tsolis, R. M., Revzin, A., Lebrilla, C. B., & Bäumler, A. J. (2017). Microbiota-activated PPAR-γ signaling inhibits dysbiotic Enterobacteriaceae expansion. Science (New York, N.Y.), 357(6351), 570–575. https://doi.org/10.1126/science.aam9949

49. Tan, B., Lu, Z., Dong, S., Zhao, G., & Kuo, M. S. (2014). Derivatization of the tricarboxylic acid intermediates with O-benzylhydroxylamine for liquid chromatography-tandem mass spectrometry detection. Analytical biochemistry, 465, 134–147.

50. Spees, A. M., Wangdi, T., Lopez, C. A., Kingsbury, D. D., Xavier, M. N., Winter, S. E., Tsolis, R. M., & Bäumler, A. J. (2013). Streptomycin-induced inflammation enhances Escherichia coli gut colonization through nitrate respiration. mBio, 4(4), e00430–13. https://doi.org/10.1128/mBio.00430-13

